# Cooperative virulence can emerge via horizontal gene transfer but is stabilized by transmission

**DOI:** 10.1101/2021.02.11.430745

**Authors:** Erik Bakkeren, Ersin Gül, Jana S. Huisman, Yves Steiger, Andrea Rocker, Wolf-Dietrich Hardt, Médéric Diard

## Abstract

Intestinal inflammation fuels *Salmonella* Typhimurium (*S*.Tm) transmission despite a fitness cost associated with the expression of virulence. Cheater mutants can emerge that profit from inflammation without enduring this cost. Intestinal virulence in *S*.Tm is therefore a cooperative trait, and its evolution a conundrum. Horizontal gene transfer (HGT) of cooperative alleles may facilitate the emergence of cooperative virulence, despite its instability. To test this hypothesis, we cloned *hilD*, coding for a master regulator of virulence, into a conjugative plasmid that is highly transferrable during intestinal colonization. We demonstrate that virulence can emerge by *hilD* transfer between avirulent strains *in vivo*. However, this was indeed unstable and *hilD* mutant cheaters arose within a few days. The timing of cheater emergence depended on the cost. We further show that stabilization of cooperative virulence in *S*.Tm is dependent on transmission dynamics, strengthened by population bottlenecks, leading cheaters to extinction and allowing cooperators to thrive.

## Main text

Bacteria often exist in dense communities. Therefore, many aspects of bacterial lifestyle are governed by social interactions^1^. This has also been observed for pathogens, which often infect hosts through collective actions^2-4^. For example, they can secrete extracellular metabolites or enzymes to assist in growth (e.g. iron-scavenging siderophores)^3^, produce toxins to compete with other species^5,6^, establish and survive within biofilms^7^, or use virulence factors to modulate the host immune response to create a favourable environment^8,9^. These collective actions function through public goods, which are costly to produce. Therefore, cheater mutants can emerge by profiting from the public good without enduring the cost of its production. In extreme cases, the overgrowth by cheaters can lead to population collapse due to the total breakdown of public good production^2^. The existence of cooperation is difficult to rationalize, although gene regulation, phenotypic heterogeneity, population structure and ecological factors^10,11^ likely contribute, if not to its emergence, at least to its stability by altering the cost/benefit ratio^2,7,12-14^.

Horizontal gene transfer (HGT) play a key role in the evolution of virulence in pathogenic bacteria^4^. In *Escherichia coli* for instance, genes encoding for secreted proteins (including virulence factors) are associated with mobile genetic elements (MGEs)^15,16^. Therefore HGT could be a mechanism by which cooperation-based virulence emerges in pathogenic bacteria^17^. However, the role of HGT in stabilizing cooperation^17^ in the long term by bringing cooperative alleles into cheaters is less clear since cheating can re-occur by mutation at the level of the vector^15^. Nevertheless, *in vitro* experiments have shown that, in structured environments, HGT increases the relatedness of neighbouring cells since proximity is required for MGE transfer. In some contexts, HGT may help to stabilize cooperation^18^. Here, we address the role of HGT in the evolution of cooperative virulence in an experimental model that captures the complexity of the host-pathogen interaction. We use a mouse model to study the emergence of cooperative virulence and its stabilization in the enteric pathogen *Salmonella enterica* serovar Typhimurium (*S*.Tm) during intestinal colonization.

*S*.Tm actively triggers gut inflammation via its Type Three Secretion System-1 (TTSS-1) to favour its own growth, but also that of related *Enterobacteriaceae* ^8,9,19^. The *Salmonella* pathogenicity island (SPI) that encodes for TTSS-1 mediated virulence (i.e., SPI-1) has been acquired by HGT^4,20,21^, as well as many other virulence determinants in *S*.Tm located on prophages and plasmids^4,22^. Expression of *ttss-1* is associated with a cost both *in vitro*^23^ and *in vivo*^9,12^, and cheaters emerge during infection^12^. The expression of *ttss-1* in *S*.Tm is bistable, allowing a subpopulation of phenotypically virulent cells to trigger inflammation, while another subpopulation grows more quickly; this division of labour strategy limits cheater out-growth^12^. However, cheaters still emerge during within-host evolution experiments^12,24^. The target of selection is the transcriptional regulation of virulence expression via HilD^12^. Such *hilD* mutants have been isolated from patients^25^ and swine^26^, and are under niche-specific positive selection according to comparative genomic analyses of more than 100 000 natural isolates of *Salmonella enterica*^27^. Therefore, division of labour alone cannot explain the maintenance of cooperative virulence in *S*.Tm and much less its emergence. We hypothesized that although HGT may facilitate the emergence of cooperative virulence, this should remain unstable. However, host-to-host transmission dynamics, expedited by population bottlenecks, should stabilize cooperative virulence, since virulent clones trigger disease and facilitate shedding whereas cheating clones do not^28^.

## Results

### Cooperative virulence can emerge *in vi*vo via HGT of the cooperative allele

We created a tractable model for the evolution of cooperative virulence by cloning *hilD* coupled to a chloramphenicol resistance cassette into the conjugative *IncI*1 plasmid P2 (aka. pCol1B9), which is native to *S*.Tm SL1344^19^. The resulting construct was named pVir. We have previously shown that P2 spreads efficiently into *S*.Tm 14028S and some *E. coli* strains *in vivo*^19,29-32^. As a donor strain, we conjugated pVir into an *S*.Tm 14028S derivative that lacks both a chromosomal copy of *hilD* and a functional *ttss*-1 locus (*invG* mutant) (**Fig. 1A**). As a recipient strain, we used a kanamycin resistant derivative of *S*.Tm 14028S that also lacks a chromosomal copy of *hilD*, but has all necessary genes to produce a functional TTSS-1 (**Fig. 1A**). Both the donor and the recipient are genetically avirulent (i.e. they do not elicit overt gut inflammation), but conjugation of pVir to the recipient should produce a transconjugant able to trigger inflammation (**Fig. 1A**). Both the donor and the recipient lack a functional TTSS-2 (*ssaV* mutant) to exclude inflammation triggered through TTSS- 2 at a later stage in mouse gut infections^33,34^.

**Figure 1.**
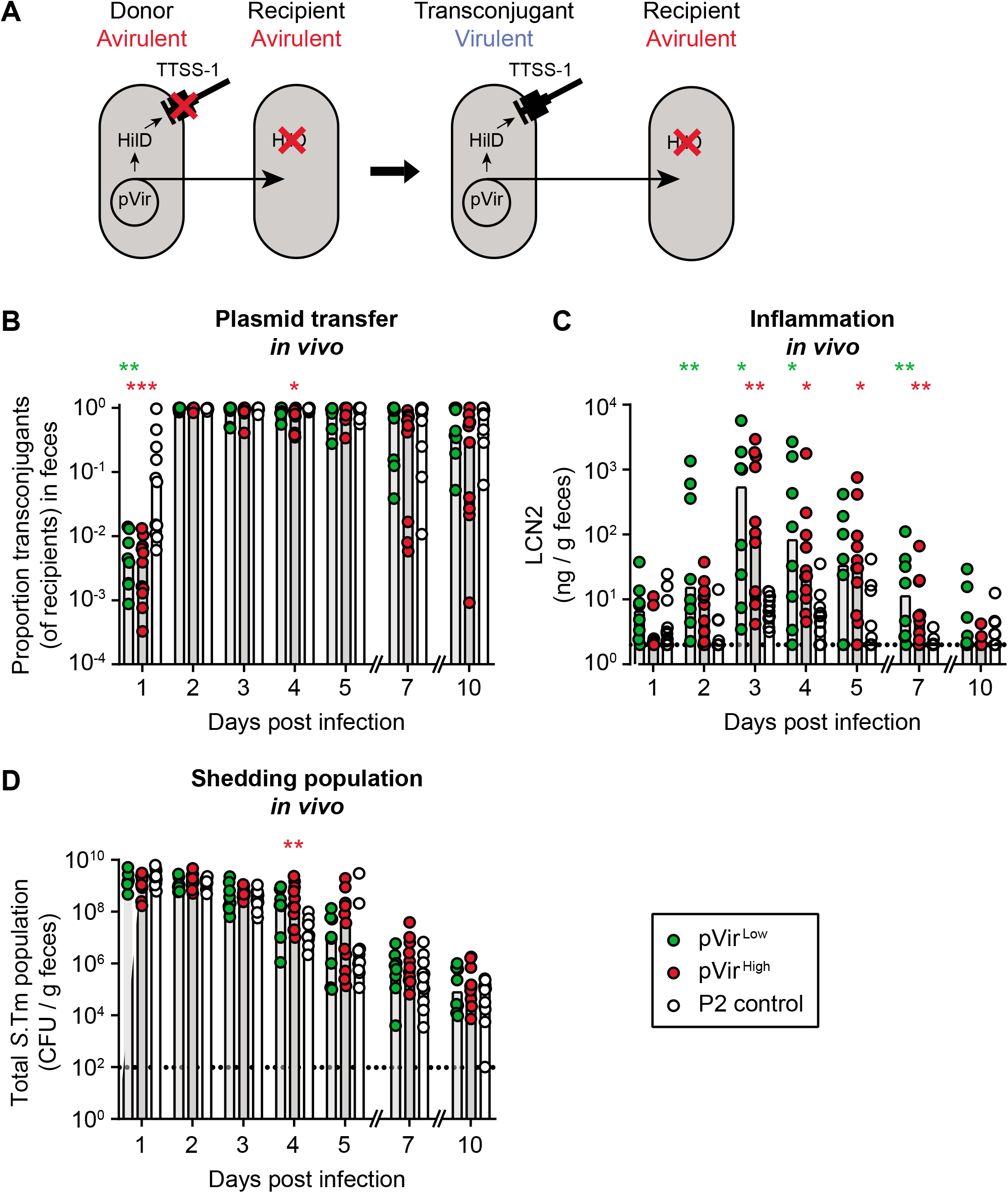
Virulence can emerge through HGT in a population of cheaters *in vivo*. **A**) Experimental system to measure maintenance of cooperative virulence by HGT. Donors contain pVir encoding *hilD*, but cannot produce a functional TTSS-1 (*invG* mutant), making them avirulent. Recipients contain all genes for a functional TTSS-1 but do not have a functional copy of *hilD* (cheaters), preventing *ttss-1* expression and virulence. Upon conjugative transfer of pVir from the donor to the recipient, a transconjugant is formed that contains both functional *ttss-1* genes and a copy of *hilD* from pVir allowing TTSS-1-mediated virulence. Transconjugants can then transfer pVir to additional recipients. **B-D**) pVir is transferred to cheater recipients and allows cooperative virulence to emerge. Ampicillin pretreated mice were sequentially infected orally with donors (14028S Δ*invG* Δ*hilD* Δ*ssaV*; Cm^R^, Amp^R^) harbouring pVir^Low^ (green; n=8), pVir^High^ (red; n=11), or P2 lacking *hilD* (control; white; n=11), and recipients (14028S Δ*hilD* Δ*ssaV*; Kan^R^, Amp^R^). Each replicate is shown and bars indicate the median. Statistics compare pVir^Low^ (green asterisks) and pVir^High^ (red asterisks) to the control on each day; Kruskal-Wallis test with Dunn’s multiple test correction (p>0.05 not significant and not indicated, p<0.05 (*), p<0.01 (**), p<0.001 (***)). Dotted line represents the detection limit. **B)** Plasmid transfer was measured by selective plating: donors Cm^R^, recipients Kan^R^, and transconjugants both Cm^R^ and Kan^R^. The proportion of transconjugants is calculated by dividing the transconjugant population by the sum of recipients and transconjugants. Replica plating was used to determine exact ratio of transconjugants compared to recipients. **C)** Inflammation was measured by a Lipocalin-2 ELISA on fecal samples. **E)** Total population was enumerated by summing all subpopulations determined by selective plating. Donor, recipient, and transconjugant populations are presented in **Fig. S3**.

To determine to which extent cooperative virulence evolution depends on the fitness cost of the cooperative allele^35^, we constructed two variants of pVir. The “low cost” variant (pVir^Low^) contains 648 bp of the regulatory region upstream of *hilD* (and all 4 transcriptional start sites characterized in^36^), while the “high cost” variant (pVir^High^) contains 279 bp of upstream regulatory region (only 2 transcriptional start sites of *hilD*^36^; **Fig. S1A**). To test the difference in cost associated with each pVir variant, we performed *in vitro* experiments comparing growth rate with *ttss-1* expression, which is costly^23^ and thus inversely correlated to growth (**Fig. S1B**-**C**). We confirmed that transconjugants harbouring pVir^Low^ expressed less *ttss-1* and grew better than those harbouring pVir^High^, confirming the difference in cost associated with *ttss-1* expression induced by these constructions (**Fig. S1B**-**C**).

To address the conditions that may support the evolution of cooperative virulence, we performed conjugation experiments in an antibiotic pretreated mouse model (modified from^37^). We introduced the donor and recipient strains sequentially into ampicillin pretreated mice at low inoculum size (10^2^ CFU donors; 10^4^ CFU recipients) to ensure that conjugation occurred only after growth in the gut. By 2 days post infection, 97% of recipients (median of all mice) obtained the plasmid, although spread of pVir^Low^ and pVir^High^ proceeded slower than the P2 control plasmid (pVir lacking *hilD*; i.e. P2cat; **Fig. 1B**, day 1 p.i.). The plasmid was also maintained in the majority of mice for the entire course of the experiment (45% of recipients; median of all mice). To test if transfer of pVir could allow the emergence of cooperative virulence in a population of avirulent recipients, we measured fecal lipocalin-2 (LCN2) as a readout for the inflammatory status of the gut. Inflammation was progressively triggered as more pVir transconjugants were formed, leading to a maximum at 3 days post infection (**Fig. 1B**-**C**). Mice containing *S*.Tm with either pVir^Low^ or pVir^High^ were significantly more inflamed than mice infected with control *S*.Tm donors (day 2-7 post infection; **Fig. 1D**). This shows that virulence can emerge within a host, since neither strain was virulent prior to conjugation **(Fig. S2)**. However, intestinal inflammation was not sustained (**Fig. 1C**), and mice began to recover leading to exclusion of *S*.Tm from the gut likely by the re-growing microbiota (**Fig. 1D; Fig. S3**)^38^. Furthermore, mice harbouring virulent *S*.Tm did not excrete significantly more *S*.Tm than control mice (**Fig. 1D; Supplementary discussion**). This observation led to two important questions: 1) why is the emergence of cooperative virulence short-lived and 2) how could virulence evolve after emergence via HGT if this is not intense or stable enough to prolong pathogen bloom?

### HGT-mediated cooperative virulence is short-lived, characterized by cost-dependent inactivation of the mobile cooperative allele

We hypothesized that the waning inflammation was a result of insufficient *ttss-1* expression. Inflammation started to decrease after day 3 post infection, while the proportion of transconjugants in the feces of mice infected with pVir-harbouring *S*.Tm remained 84% at day 4 post infection (median of all mice with pVir^High^ or pVir^Low^; **Fig. 1B**). Therefore, plasmid loss could not explain cooperative virulence loss. However, cheating could have occurred by mutations on the plasmid, as previously suggested^15,18^.

To address this, we performed a Western blot on transconjugant colonies isolated from feces (“colony blot”)^12,24^ to probe for *ttss-1* expression. As expected, cooperative virulence emergence by HGT was transient, since clones that do not express *ttss-1* arose (**Fig. 2A**). We performed whole-genome sequencing on evolved clones that were either *ttss-1*^+^ or *ttss-1*^-^ as determined by the colony blot. It showed that cheating was a result of mutations or deletions in either the coding sequence or regulatory regions of *hilD* on pVir, and not due to chromosomal changes (**Tables S1-4**). This further supported the hypothesis that maintaining cooperation by HGT is inefficient, since cheating now occurred at the level of the MGE. Strikingly, the loss of *ttss-1*^+^ clones was slower with pVir^Low^ compared to pVir^High^, leading to a higher proportion of cooperating clones bearing pVir^Low^ by the end of the experiment (**Fig. 2A**). This indicates that cost influences the maintenance of cooperative virulence mediated by HGT (as predicted by theory^15,17^), extending our previous *in vivo* work on the cost-dependent cheating dynamics of *S*.Tm with the chromosomal copy of *hilD*^12^.

**Figure 2.**
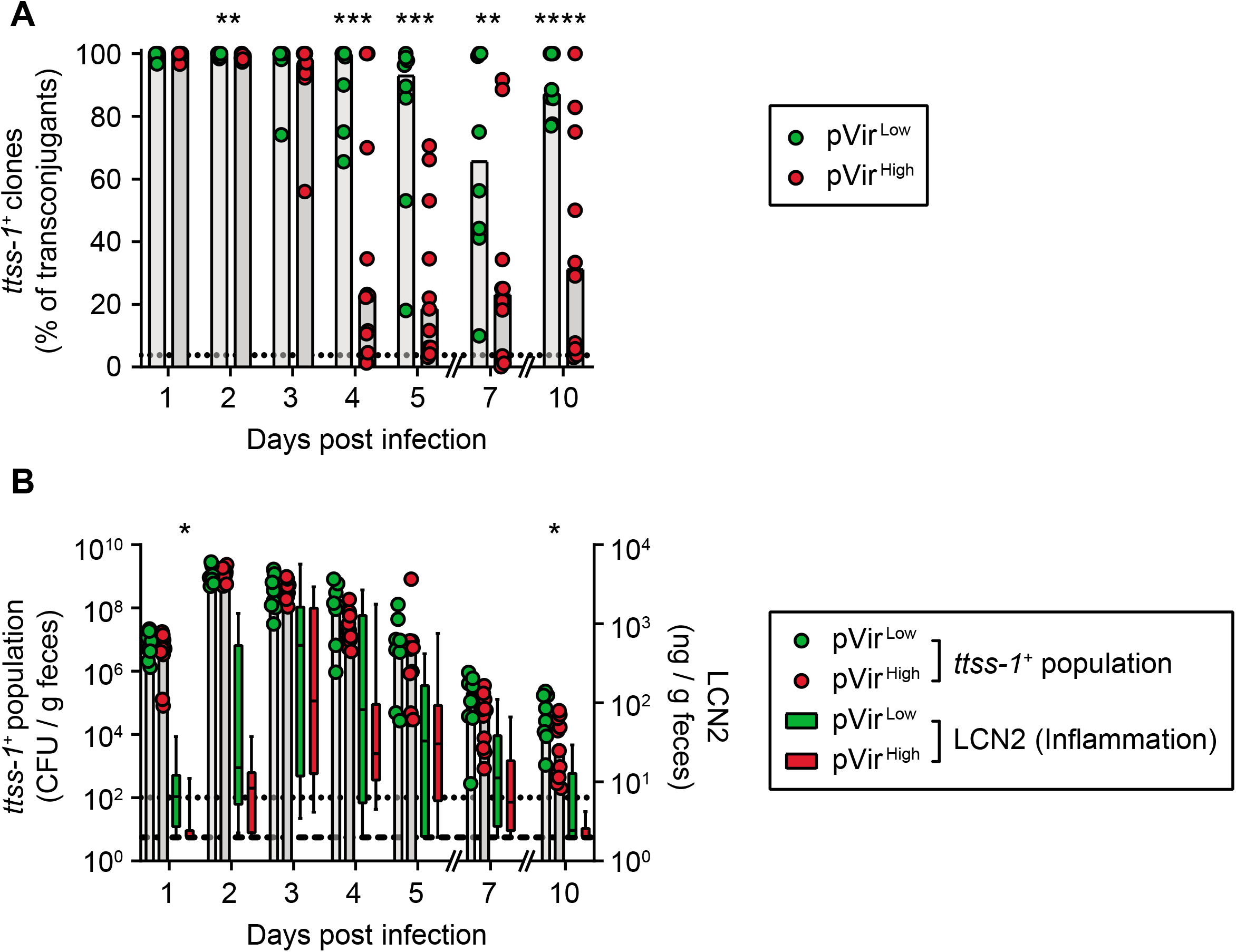
The restoration of cooperative virulence by HGT is short-lived and depends on fitness cost. The proportion of cooperators in the feces of mice in **Fig. 1D**-**F** was analyzed by colony western-blot. Transconjugants carrying pVir^Low^ (green) and pVir^High^ (red) were analyzed and compared using a two-tailed Mann-Whitney U test (p>0.05 not significant and not indicated, p<0.05 (*), p<0.01 (**), p<0.001 (***), p<0.0001 (****)). **A)** Transconjugants lose the ability to express *ttss-1*. MacConkey plates containing colonies of transconjugants were analyzed for expression of SipC as a proxy for *ttss- 1* expression; the percentage of colonies that expressed SipC are reported out of the total transconjugant population. Each data point is shown and bars indicate the median. The black dotted line indicates the conservative detection limit for the colony blot, which is dependent on the number of colonies on the plate (values can therefore appear below the detection limit). **B)** The size of the population able to express *ttss-1* correlates with inflammatory state of the mouse. The transconjugant population size was multiplied by the proportion of cooperating clones (from **panel A)** to determine the size of the population able to express *ttss-1* (*ttss-1*^+^ population; plotted on the left y-axis; all data points plotted and line indicates the median). The black dotted line indicates the detection limit from selective plating. The inflammatory state is shown as a box (median indicated with a line and the quartiles are defined by the edges of the box) and whiskers define the minimum and maximum values (same data as in **Fig. 1C**). The dashed black line indicates the detection limit.

However, since both the cheating dynamics and the total population size dictate the size of the population able to trigger inflammation, we multiplied the proportion of transconjugants able to trigger inflammation (from **Fig. 2A**) with the transconjugant population size (**Fig. S3**) to obtain the effective size of the cooperative population (i.e., the *ttss-1*^+^ population; **Fig. 2B**). We observed a rise in the cooperative population due to plasmid transfer correlating with the onset of inflammation (day 1-2 p.i., **Fig. 2B**), followed by a drop in the cooperative population associated with the waning inflammation between days 5-10 p.i., **Fig. 2B**). This supported the hypothesis that the loss of inflammation could at least be partly driven by the loss of the cooperative population. As predicted by analyzing the proportion of transconjugants able to express *ttss-1* (**Fig. 2A**), the cooperative population was higher in mice infected with pVir^Low^ harboring cells compared to those infected with pVir^High^ harboring cells at the end of the experiment (Day 10; **Fig. 2B**).

Next, we addressed if HGT could help in maintaining cooperation in a population of recently emerged cooperators competing against remaining cheaters. In this experiment, the donor contained a functional *invG* allele, making the donor virulent (in comparison to the scenario in **Fig. 1** where the donor is avirulent). We performed a competitive infection between a *hilD* mutant (i.e., cheater) and the donor containing pVir^Low^ (i.e., cooperator) at an equal ratio (low inoculum size; ∼10^2^ CFU of each strain, introduced sequentially to avoid plasmid transfer in the inoculum). Importantly, we performed this experiment in two configurations (**Fig. 3A**): in the first group which we called “mobile pVir”, we used the same recipient cheater strain as in **Fig. 1**, which can obtain pVir from the cooperator; in the second group called “non-mobile pVir”, we used a cheater strain that carried P2 (P2 does not confer a fitness advantage to competing 14028S strains^19,32^) labelled with a kanamycin resistance marker. In this case, since pVir and P2 are incompatible and have mechanisms for entry exclusion^39^, plasmid transfer cannot be detected (confirmed by selective plating). As expected, when pVir was mobile, the plasmid was maintained in the population for longer compared to the non-mobile scenario, in which the cooperating strain was outcompeted by the cheating strain (**Fig. 3B-C; Fig. S4**). Furthermore, inflammation was maintained for longer in mice with the mobile pVir scenario (**Fig. 3D**), which was reflected in a trend towards higher shedding populations (**Fig. 3E**). Importantly, in some mice with the mobile pVir scenario, the inflammation and the shedding population also diminished over time (**Fig. 3D-E**). Therefore, we measured the proportion of *ttss-1*-expressing clones in the population at the end of the experiment. Although more *ttss-1*-expressing clones were observed in the mobile pVir scenario compared to when pVir was not mobile (**Fig. 3F**), clones that did not express *ttss-1* were detected within the pVir-containing population (**Fig. 3G**).

**Figure 3.**
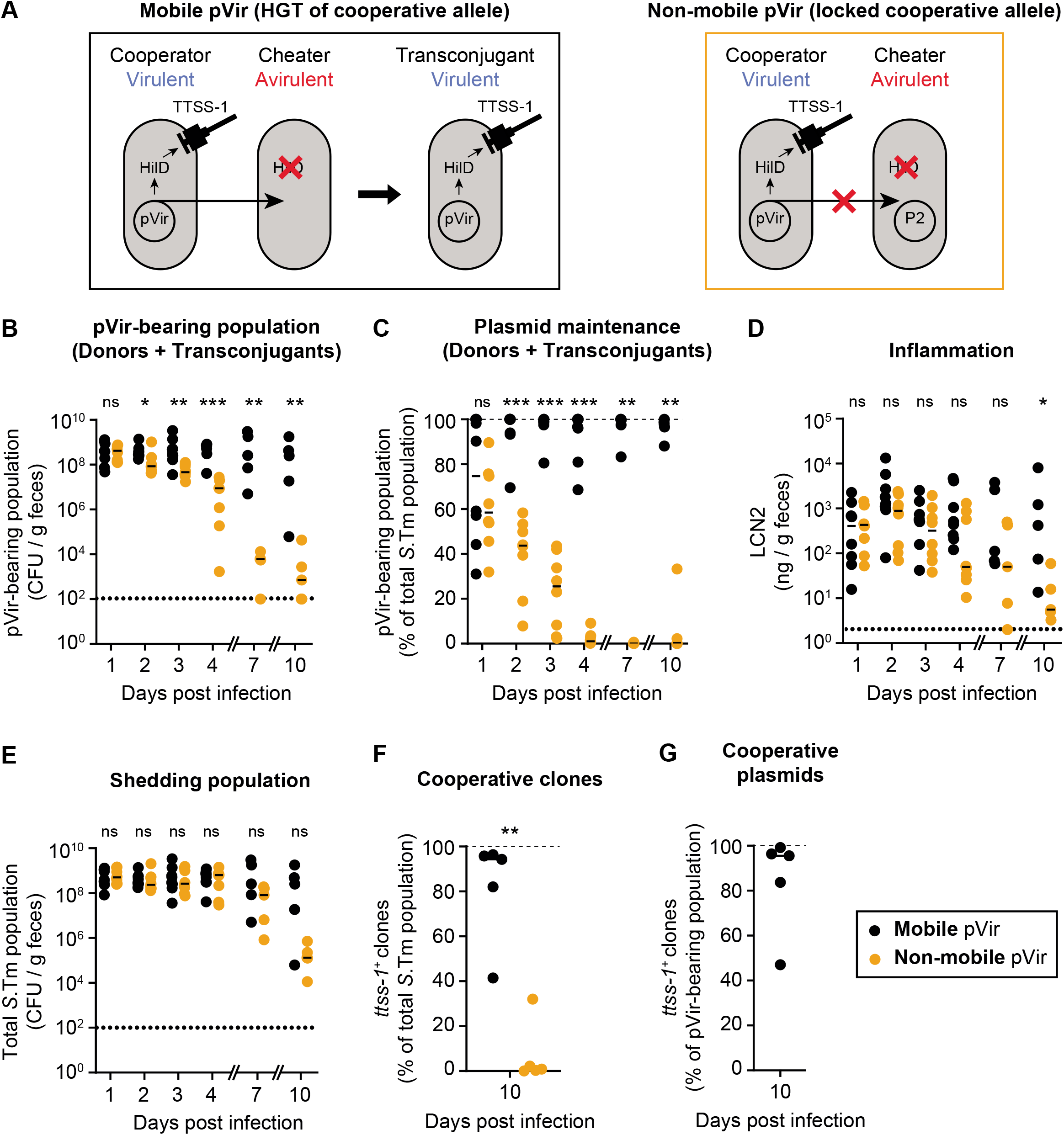
HGT can increase the duration of, but does not stabilize, cooperative virulence. **A)** Experimental system to determine the role of HGT in the stabilization of cooperative virulence. In both the mobile pVir and non-mobile pVir scenarios, the cooperator contains pVir^Low^ and a functional TTSS-1, making it virulent. In the mobile scenario, pVir can be transferred to cheaters. In the non- mobile scenario, transfer of pVir is blocked because of an incompatible plasmid, P2, in the cheater strain. **B-G)** Ampicillin pretreated mice were orally infected with ∼10^2^ CFU of cheater (14028 Δ*hilD* Δ*ssaV*; Kan^R^, Amp^R^) immediately followed by ∼10^2^ CFU of pVir^Low^ donor (14028 Δ*hilD* Δ*ssaV* pVir; Cm^R^ encoded on pVir, Amp^R^). The donor contained a functional *invG* allele, making it virulent (*ssaV* is deleted in both strains). Mice (n=8 until day 4; n=5 until day 10 for both groups) were given either a cheater with no plasmid (i.e., same strain used in **Fig. 1D-F**; <u>mobile pVir</u>; black) or a cheater with P2 (incompatible with pVir; <u>non-mobile pVir</u>; orange). Dotted lines indicate detection limits. Medians are indicated by lines. Two-tailed Mann-Whitney U tests (p>0.05 (ns), p<0.05 (*), p<0.01 (**), p<0.001 (***)) are used to compare the mobile pVir and non-mobile pVir scenarios on each day. Donor, recipient, and transconjugant populations are presented in **Fig. S4. B**) The pVir-bearing population was determined by selective plating on Cm-supplemented MacConkey agar. **C)** the pVir- bearing population is reported as a percentage of the total population. Dashed line indicates 100% plasmid spread. **D)** Inflammation was measured by a Lipocalin-2 ELISA on fecal samples. **E)** Total *S*.Tm populations determined by the sum of Cm- and Kan-supplemented MacConkey agar. **F-G)** Cm- supplemented MacConkey agar plates containing colonies from feces collected on day 10 post infection were analyzed for SipC expression as a proxy for *ttss-1* expression using a colony Western blot and represented as the percentage of colonies that expressed SipC are reported out of the total *S*.Tm population **(F)** or out of the pVir-containing population for the mobile pVir scenario **(G)**.

Altogether, this indicates that HGT can increase the time that cooperating clones exist, but that HGT alone cannot stabilize cooperation within a host. This is in line with theory and *in vitro* work in well-mixed environments where cheaters benefit from public good produced by cooperators^15,18^.

### Disease in new hosts depends on the proportion of transmitted cooperators

In our model system, cooperative virulence can emerge via HGT, however it remains unstable within-host. In the case of *S*.Tm, TTSS-1-triggered inflammation has two important consequences that can nevertheless promote cooperative virulence: it favors transmission of *S*.Tm^8,40^ and fosters pathogen blooms in the next host. Therefore, HGT may influence the evolution of *S*.Tm by increasing the duration of shedding of virulent clones able to trigger inflammation in a new host. Selection for cooperative virulence should be a result of increased benefit after transmission.

To address this hypothesis, we took fecal suspensions from mice in **Fig. 1** on day 2 p.i. (the maximum population size of *ttss-1*-expressing clones; **Fig. 2B**) and day 10 p.i. (the minimum population size of *ttss-1*-expressing clones; **Fig. 2B**) and transferred them into new ampicillin pretreated mice (**Fig. 4A**).

**Figure 4.**
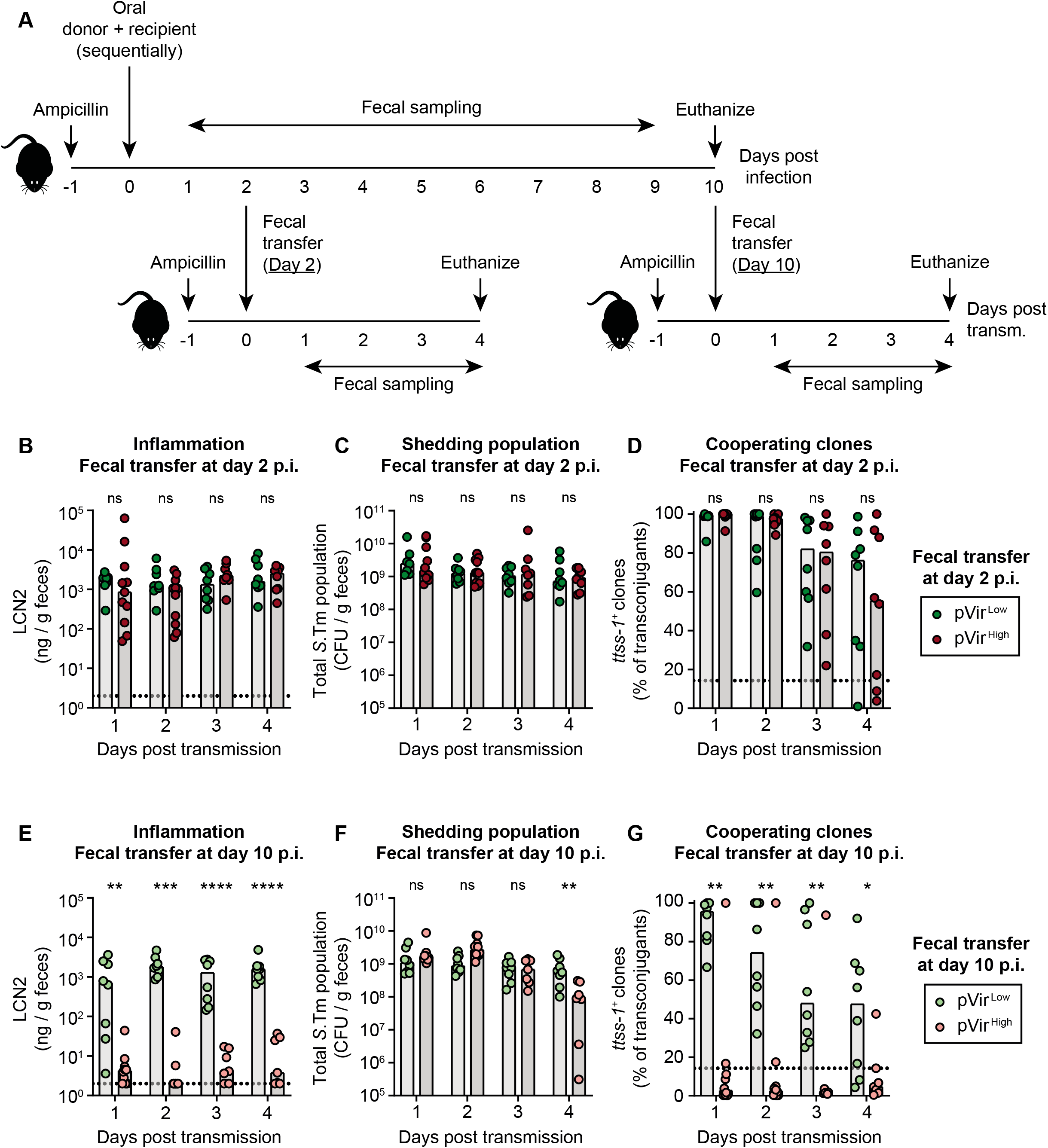
Successful infections in new hosts depend on the proportion of transmitted cooperators. **A)** Experimental scheme for transmission experiments. Feces from mice in **Fig. 1B-D** collected on day 2 and day 10 post infection were suspended in PBS and given to new ampicillin pretreated mice. **B-G)** Mice orally given fecal resuspensions with *S*.Tm harbouring pVir^Low^ (green; dark shade for day 2 transmission; light shade for day 10 transmission; n=8) are compared to pVir^High^ (red; dark shade for day 2 transmission; light shade for day 10 transmission; n=11) using a two-tailed Mann-Whitney U test (p>0.05 (ns), p<0.05 (*), p<0.01 (**), p<0.001 (***), p<0.0001 (****)). All data points are shown and medians are represented by bars. **B-D)** Mice given feces from day 2 post infection. **E-G)** Mice given feces from day 10 post infection. **B**,**E)** Inflammation was quantified using a LCN2 ELISA. The dotted lines indicate the detection limit. **C**,**F)** The shedding population was enumerated by summing all populations determined by selective plating. Donor, recipient, and transconjugant populations are presented in **Fig. S2. D**,**G)** MacConkey plates containing colonies of transconjugants were analyzed for expression of SipC as a proxy for *ttss-1* expression using a colony Western blot; the percentage of colonies that expressed SipC are reported out of the total transconjugant population. The black dotted line indicates the conservative detection limit for the colony blot, which is dependent on the number of colonies on the plate (values can therefore appear below the detection limit).

When fecal populations taken from day 2 p.i. were transferred into new mice, inflammation was triggered, and there were no significant differences between mice infected with pVir^Low^ or pVir^High^ *S*.Tm carriers (**Fig. 4B**). This led to consistent shedding over the course of the experiment in all the recipient mice (**Fig. 4C; Fig. S5A-C**). This is likely attributed to the high proportion of cooperating clones transferred with samples from day 2 p.i. (**Fig. 2**). However, in all mice, cheaters arose (**Fig. 4D**), further supporting the instability of cooperative virulence in this system. In contrast, when feces from day 10 p.i. were transferred, only the recipient mice infected with *S*.Tm harbouring pVir^Low^ became inflamed (**Fig. 4E**). This was reflected in the shedding population at day 4 post transmission, where mice infected with *S*.Tm harbouring pVir^Low^ contained significantly more *S*.Tm in the feces compared to mice infected with *S*.Tm containing pVir^High^ (**Fig. 4F; Fig. S5D-F**). These differences were likely a result of the proportion of cooperative clones in the feces of the donor mice at day 10 p.i.: mice infected with *S*.Tm pVir^Low^ had significantly more cooperators than mice infected with *S*.Tm pVir^High^ (**Fig. 2**). Moreover, as in the fecal transfer at day 2 p.i., in all mice, cheaters arose and outgrew cooperators (**Fig. 4G**). Interestingly, three mice with *S*.Tm pVir^High^ that contained a high proportion of cooperative clones at day 10 p.i. (**Fig. 2A**) did not lead to strong inflammation after transmission (**Fig. 4E**). This could indicate that additional cost-dependent factors influence the ability to trigger inflammation after transmission. Nevertheless, since transmission of feces containing pVir^High^ *S*.Tm led to inflammation dependent on the proportion of cooperators (e.g. compare the fecal transfer at day 2 p.i. (**Fig. 4B**) to the transfer at day 10 p.i. (**Fig. 4E**)), we concluded that the proportion of cooperators do contribute to transmission dynamics.

Altogether, this indicates that the proportion of cooperators, which is influenced by the cost, has implications in triggering disease and prolonging gut luminal growth in the next host after transmission. However, since cheaters can also be transmitted in the presence of enough cooperators, this can lead to a progressive accumulation of cheater strains^17^ (compare **Fig. 2A to Fig. 4D**,**G**) indicating that, under these conditions, this process cannot be supported indefinitely.

### Narrow transmission bottlenecks promote the stability of cooperative virulence

It has been proposed that cooperation can be stabilized by repetitive population bottlenecks, that is, when few founding members lead to the establishment of new populations^11^. The same process could apply to *S*.Tm cooperative virulence as the population is subject to population bottlenecks during host-to-host transmission. Bottlenecks can be the result of environmental stress between hosts, dilution during transmission or due to competition against the resident microbiota before establishing a favorable niche in the gut^41^. To address this experimentally, we simulated a narrow population bottleneck by infecting new mice with single evolved clones from mice in **Fig. 1**. We used clones evolved in different mice, representing both cooperators (i.e., *ttss-1*^+^ clones) and cheaters (i.e., *ttss-1*^-^ clones) in both the high cost and low cost variants (3 clones per group; 2-3 new mice infected per clone). For both pVir^Low^ and pVir^High^, *ttss-1*^+^ clones were able to trigger inflammation and *ttss-1*^-^ clones were not (**Fig. 5A**). Again, this was reflected in the shedding population, where mice infected with *ttss-1*^+^ clones shed significantly more *S*.Tm on day 4 p.i. compared to mice infected with *ttss-1*^-^ clones (**Fig. 5B**). Note that the antibiotic pretreatment allows pure cheater populations to reach the same population size as cooperators for three days post- infection.

**Figure 5.**
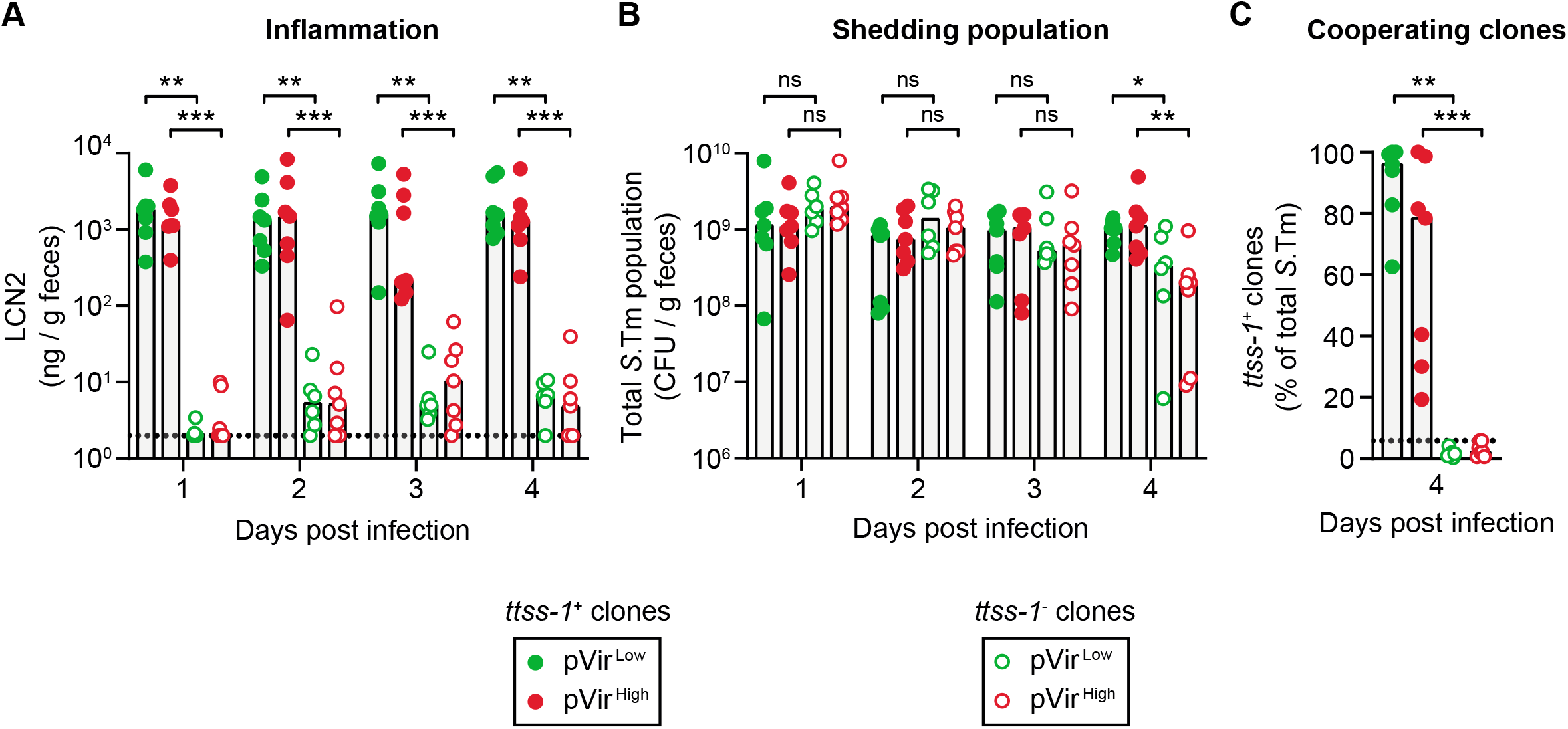
Evolved cooperating clones can trigger inflammation and lead to shedding, while evolved cheating clones cannot. Ampicillin pretreated mice were orally infected with evolved transconjugant clones isolated from day 7 or day 10 from mice in **Fig. 1B**-**D**. 3 cooperating clones (solid circles; *ttss-1*^+^ clones) and cheating clones (hollow circles; *ttss-1*^*-*^ clones) were randomly chosen for each of pVir^Low^ (green) or pVir^High^ (red). Each clone was infected into 2-3 mice (∼5×10^7^ CFU inoculum), leading to a total of 6-7 mice per group. All clones were whole-genome sequenced (mutations and indels are summarized in **Tables S1-4**): *ttss-1*^+^ pVir^Low^ (Z2296 (3 mice), Z2306 (2 mice), Z2310 (2 mice); n=7), *ttss-1*^+^ pVir^High^ (Z2238 (3 mice), Z2246 (2 mice), Z2253 (2 mice); n=7), *ttss-1*^-^ pVir^Low^ (Z2298, Z2301, Z2305; 2 mice per clone; n=6), *ttss-1*^-^ pVir^High^ (Z2239 (3 mice), Z2243 (2 mice), Z2311 (2 mice); n=7). All data points are shown and medians are indicated by bars. Comparisons are made between *ttss-1*^+^ and *ttss-1*^-^ clones (for each of pVir^Low^ and pVir^High^) using a two-tailed Mann-Whitney U test (p>0.05 (ns), p<0.05 (*), p<0.01 (**), p<0.001 (***)). **A)** Inflammation was quantified using a LCN2 ELISA. The dotted lines indicate the detection limit. **B)** The shedding population was enumerated on MacConkey agar. **C)** MacConkey plates containing colonies were analyzed for expression of SipC as a proxy for *ttss-1* expression using a colony western blot; the percentage of colonies that expressed SipC are reported out of the total transconjugant population. The black dotted line indicates the conservative detection limit for the colony blot, which is dependent on the number of colonies on the plate (values can therefore appear below the detection limit).

As a control, we measured the proportion of cooperative clones in these mice on day 4 p.i. As expected, mice infected with cheater clones contained only *ttss-1*^-^ clones and mice infected with cooperative clones contained mostly *ttss-1*^+^ clones, although cheaters began to emerge in these mice as well (**Fig. 5C**).

Importantly, the proportion of cooperators in mice infected with single *ttss-1*^+^ clones appeared higher than in mice that received a non-isogenic mixture of cooperators and cheaters (compare **Fig. 5C** to **Fig. 4D**,**G**). This suggests that population bottlenecks could indeed promote cooperation, since these processes increase the probability of monoclonal infections. To support this prediction, we tested the correlation between the proportion of cooperators given to mice in the transmission experiments (**Fig. 4** and **Fig. 5**) and inflammation, the shedding population, and the proportion of cooperators at day 4 post infection (**Fig. S6**). The proportion of cooperators in the input correlated with the resulting inflammation, shedding, and the final proportion of cooperative clones (**Fig. S6**; slopes are significantly non-zero: **Fig. S6A** p<0.0001; **Fig. S6B** p=0.0002; **Fig. S6C** p<0.0001), in line with previous work comparing the proportion of the population able to express the TTSS-1^+^ phenotype and the resulting inflammation^9^.

## Discussion

Our results provide the first comprehensive analysis of different aspects that could influence cooperative virulence in an infection model. We show that although cooperative virulence can emerge in *S*.Tm from two initially avirulent strains by HGT, the role of HGT was limited to the emergence and transient maintenance of cooperative virulence, but not its evolutionary stability (**Fig. 1, 2 and 3**). However, transmission dynamics (i.e., timing and bottlenecks) are essential to sustain cooperative virulence (**Fig. 4, 5**). Indeed, we observed that cheating re-occurred in transconjugants via inactivation of the cooperative allele by mutation (**Table S1-S4**). Because both the cooperative and cheating alleles on MGEs spread equally well in the population, HGT only transiently maintains cooperation within a population. This is in line with previous studies in well-mixed environments *in vitro* and in *in silico* models^15,18,42^. Moreover, higher fitness cost in our experimental system led to greater instability of cooperative virulence (**Fig. 2, 4**), which further highlights that the success of evolution of cooperation by HGT is dictated by the cost of the cooperative allele^17,35^. Complex fine-tuned regulation is essential to ensure virulence stability in *S*.Tm^12^. Accordingly, *hilD* is integrated into the chromosome of *S*.Tm within SPI-1, which, compared to carriage on multi-copy plasmids, could favor the evolution of a regulation deeply entangled with the general physiology of the pathogen^43-45^.

Theoretical work proposes that transmission likely plays a role in the epidemiological success of virulence^17,46^, and that population bottlenecks influence this process^11^. This is confirmed in our experiments, where we show that a high proportion of cooperating clones trigger disease and lead to greater shedding than when a low proportion of cooperative clones reached a new host, for instance when transmission occurs late (**Fig. 4, 5, S6**). This is because cheaters alone cannot free-ride off of the inflammation normally triggered by cooperators (**Fig. S7**). In the case of *S*.Tm, population fragmentation occurs when a subset of the population is released from a host through fecal pellets, and bottlenecks occurs in harsh external environment or during colonization of a recipient host in which colonization resistance is mediated by the protective gut microbiota. Therefore, environmental factors such as diet perturbations^29,47^ or exposure to antibiotics^37^ can widen transmission bottleneck and have profound implications on the evolution of virulence.

Overall, we suggest that blooming after successful transmission is the primary selective force to stabilize cooperative virulence in a host infection model. *S*.Tm is a case where the benefit of cooperative virulence directly fuels its transmission^8,9,12,40^, making transmission-dependent selection fairly intuitive. The same could apply for other enteric pathogens such as *Vibrio cholerae* (encoding the cholera toxin on a phage)^48^, *Shigella* spp. (containing phage- and plasmid-encoded secreted virulence factors)^49^, or *Yersinia* spp. (encoding the Yop virulon on a plasmid)^50^, which all use secreted virulence factors to survive in the host and/or directly increase shedding. However, the general selection for virulence determinants with extracellular functions could also be driven by transmission-dependent bottlenecks that increase relatedness, excluding cheaters and favouring cooperators^35^. The only requirement for this process would be that a given virulence factor increases the probability for the pathogen to survive in the next host when more virulence factor-encoding cells are present. This could apply, for example, to siderophore production or quorum sensing systems. Further work is required to explore the generality of transmission-dependent stabilization of cooperative virulence.

Studies that investigate bacterial sociality and the evolution of virulence such as the one we present are critical given the current antibiotic crisis^51^, as they aid the discovery of antibiotic-free treatments to manage bacterial infections. Research into the social aspects of infection such as collective virulence factor secretion has inspired “anti-virulence compounds” (that target extracellular virulence factors) as a therapeutic avenue for minimizing resistance^52^. Additionally, exploiting cheating behaviour to destabilize cooperation has been suggested as a possible therapeutic strategy (coined “Hamiltonian medicine”)^53^. In extension of our previous work^12,28^ on *S*.Tm, and as suggested theoretically^54^ and experimentally with *P. aeruginosa*^55^, we show here that administering *hilD* mutants has the potential to destabilize virulence-mediated *Salmonella* blooms and transmission. Furthermore, strategies that slow down HGT, such as vaccination^30-32^, can be useful for both reducing antibiotic resistance plasmid spread but also reducing the emergence or maintenance of virulence in populations.

## Materials and methods

### Strains, plasmids, and primers used in this study

All the strains and plasmids used in this work are summarized in **Table 1** and **Table 2**. Bacteria were grown in lysogeny broth (LB) media containing the appropriate antibiotics (50 µg/ml streptomycin (AppliChem); 15 µg/ml chloramphenicol (AppliChem); 50 µg/ml kanamycin (AppliChem); 100 µg/ml ampicillin (AppliChem)) at 37°C (or 30°C if containing pKD46 or pCP20). Gene deletion mutants were performed using the λ *red* system^56^. Desired genetic constructs were transferred into the appropriate background strain using P22 HT105/1 *int-201* phage transduction^57^. Antibiotic resistance cassettes were removed using the heat inducible FLP recombinase encoded on pCP20, if desired^56^. Expression vectors (e.g. pM975 and pM972) were transformed into the desired strain using electroporation.

**Table 1.**
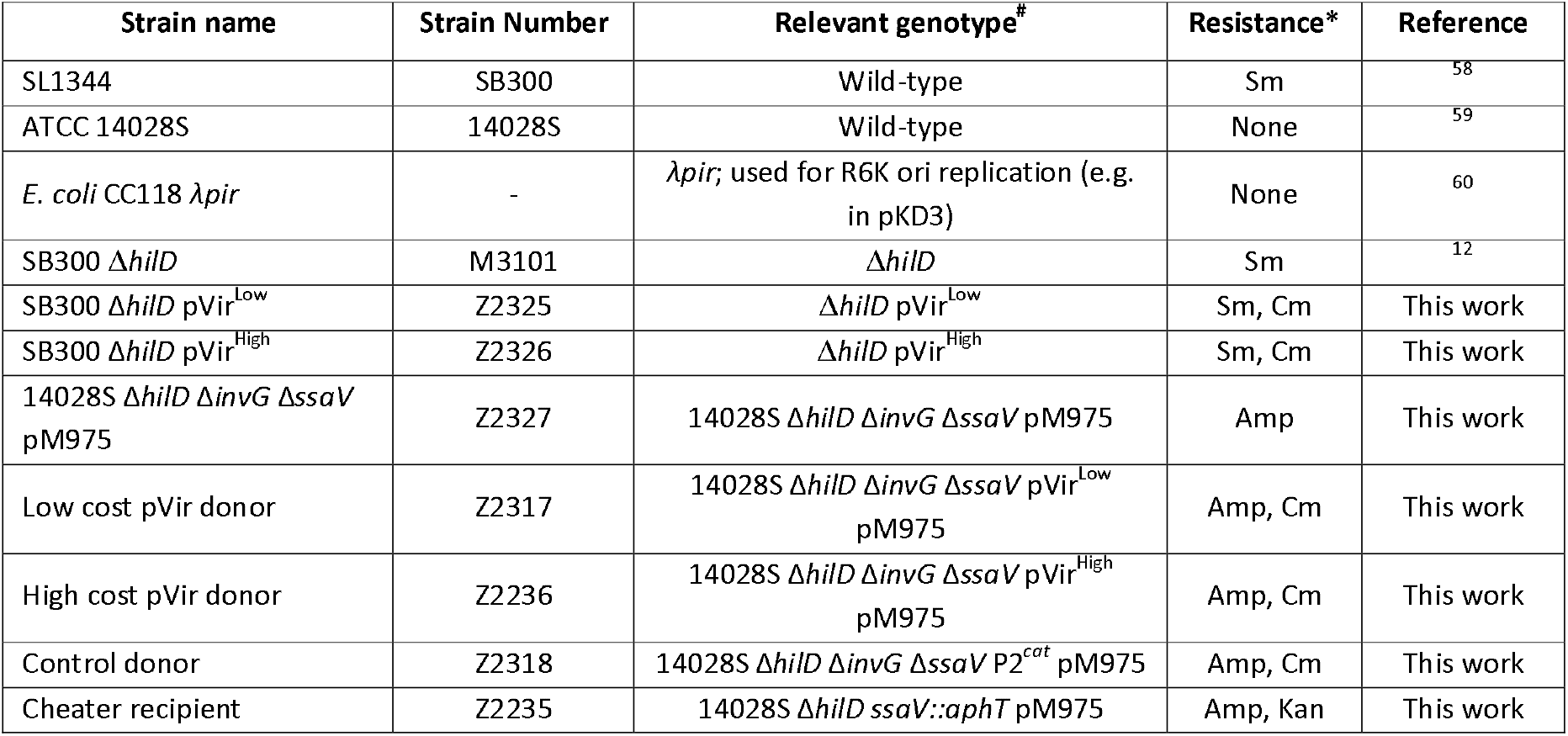
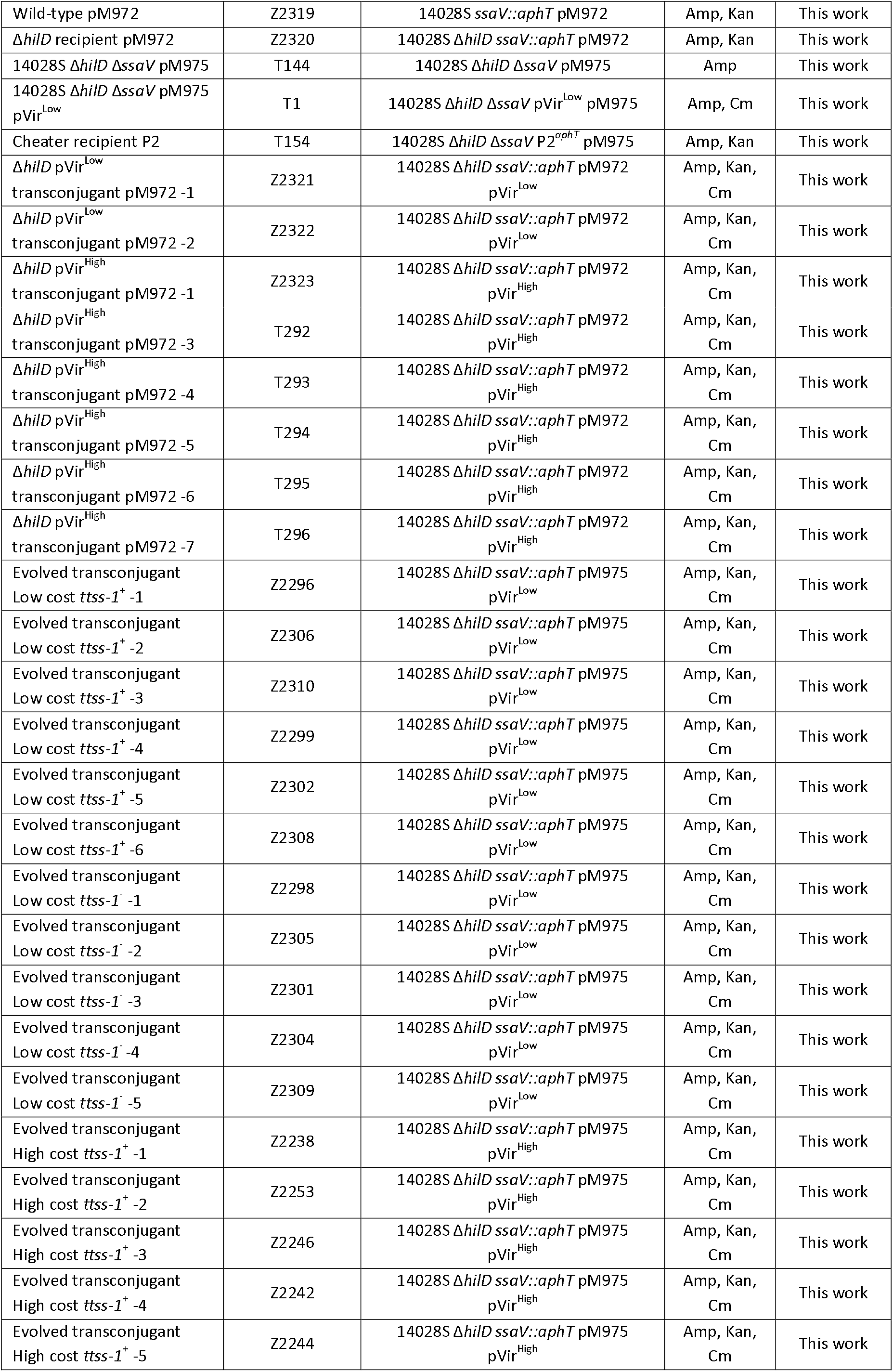
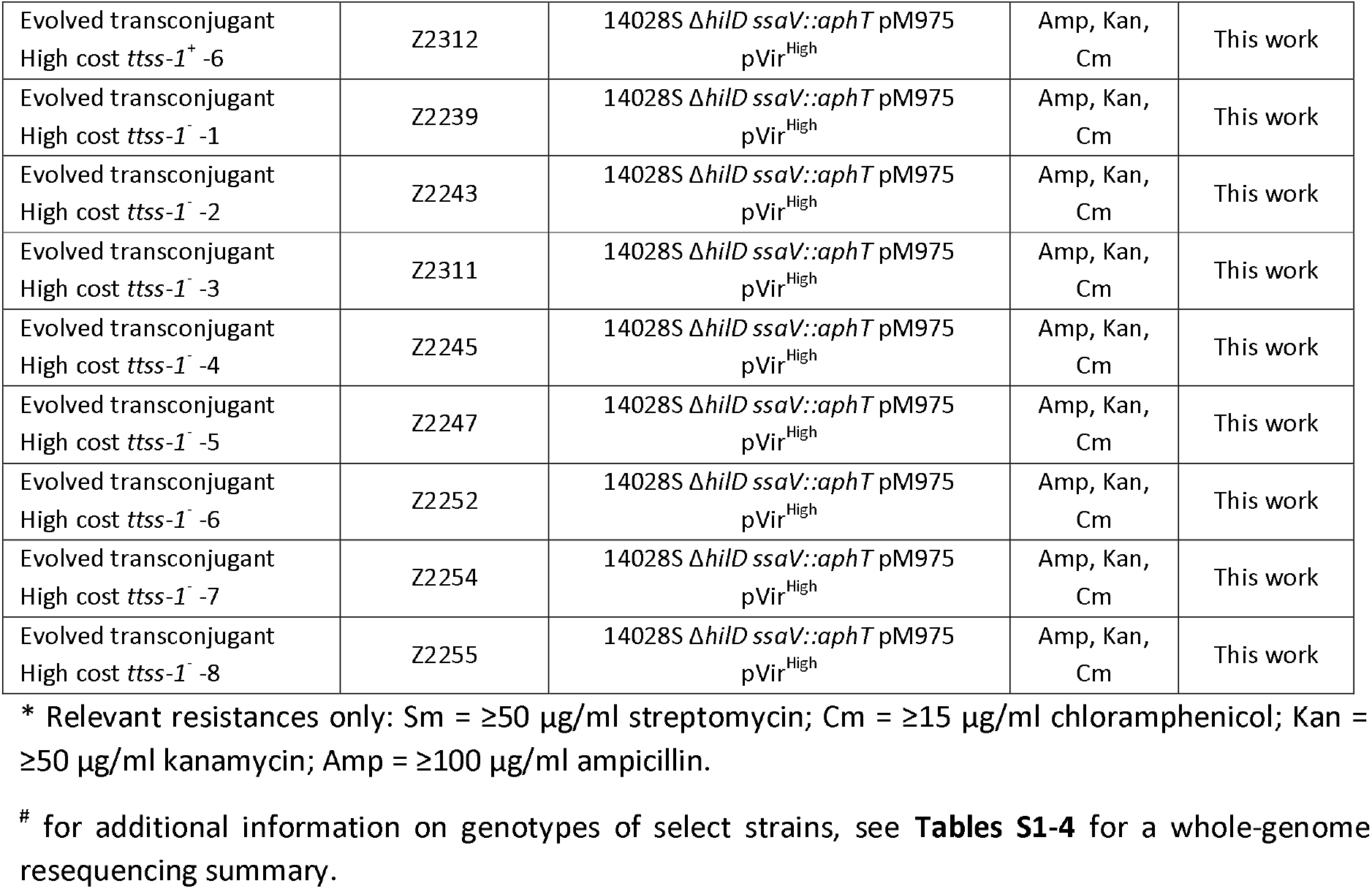
Strains used in this study.

**Table 2.**
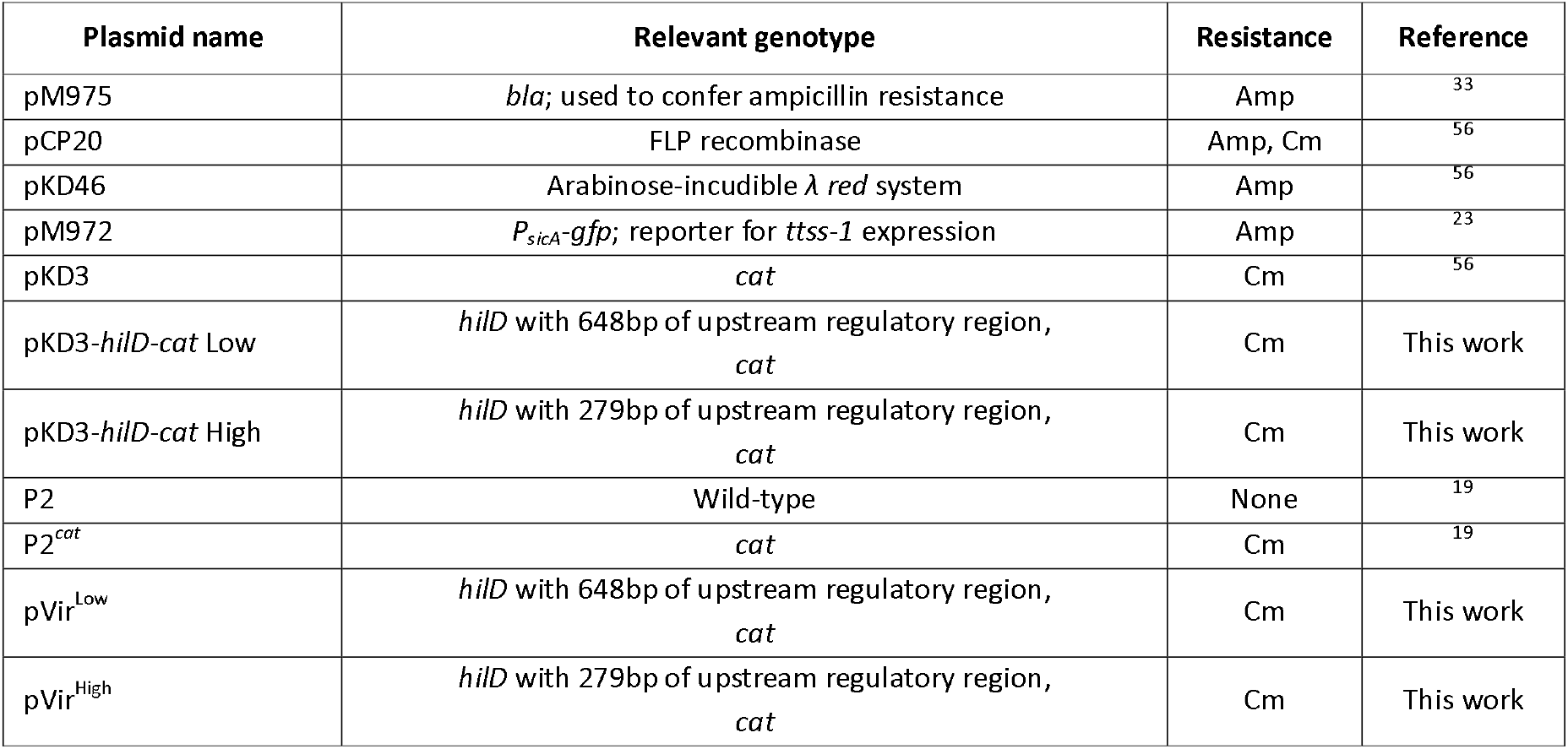
Plasmids used in this study.

To create pVir donor strains, *hilD* was amplified with PCR using high-fidelity Phusion polymerase (ThermoFisher Scientific) from the chromosome of SL1344 with either 648bp (low cost) or 279bp (high cost) of regulatory region (**Fig. S1**) and cloned into pKD3 upstream of the chloramphenicol resistance cassette using Gibson Assembly (NEB). Primers to amplify *hilD* contained ∼40bp homology to the sites flanking a *Nde*I site in pKD3. pKD3 was digested with *Nde*I, purified, and mixed with the PCR amplicon in Gibson Assembly Master Mix (NEB; protocol as described by the manufacturer). The products were transformed into *E. coli* CC118 λ*pir*, and colonies were verified to contain the desired plasmid through PCR and Sanger sequencing. The resulting *hilD-cat* construct was then amplified from cloned plasmid with Phusion PCR using primers with homology to the target site in P2 (upstream of the colicin Ib locus; *cib*) and introduced into SB300 Δ*hilD* using λ *red*^*56*^. Positive clones were determined by PCR, leading to pVir^Low^ and pVir^High^. Lastly, the pVir plasmids were conjugated *in vitro* into the desired strain by mixing the 10^5^ CFU from an overnight culture of the donor strain with the desired recipient, allowing conjugation overnight at 37°C on a rotating wheel, and plating the cells on MacConkey agar to select for transconjugants. For *in vivo* experiments, pVir plasmids were conjugated into 14028S Δ*hilD* Δ*invG* Δ*ssaV* pM975 or 14028S Δ*hilD* Δ*ssaV* pM975. For *ttss-1* expression analysis, pVir plasmids were conjugated into 14028S Δ*hilD ssaV::aphT* pM972. All primers used for strain or plasmid construction and verification are listed in **Table 3**.

**Table 3.**
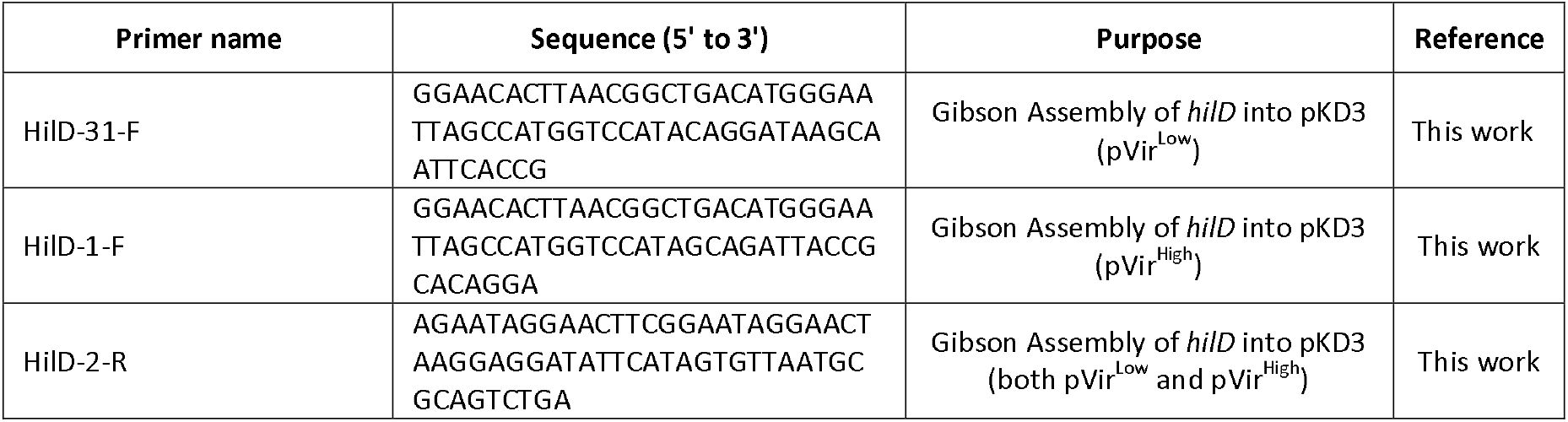
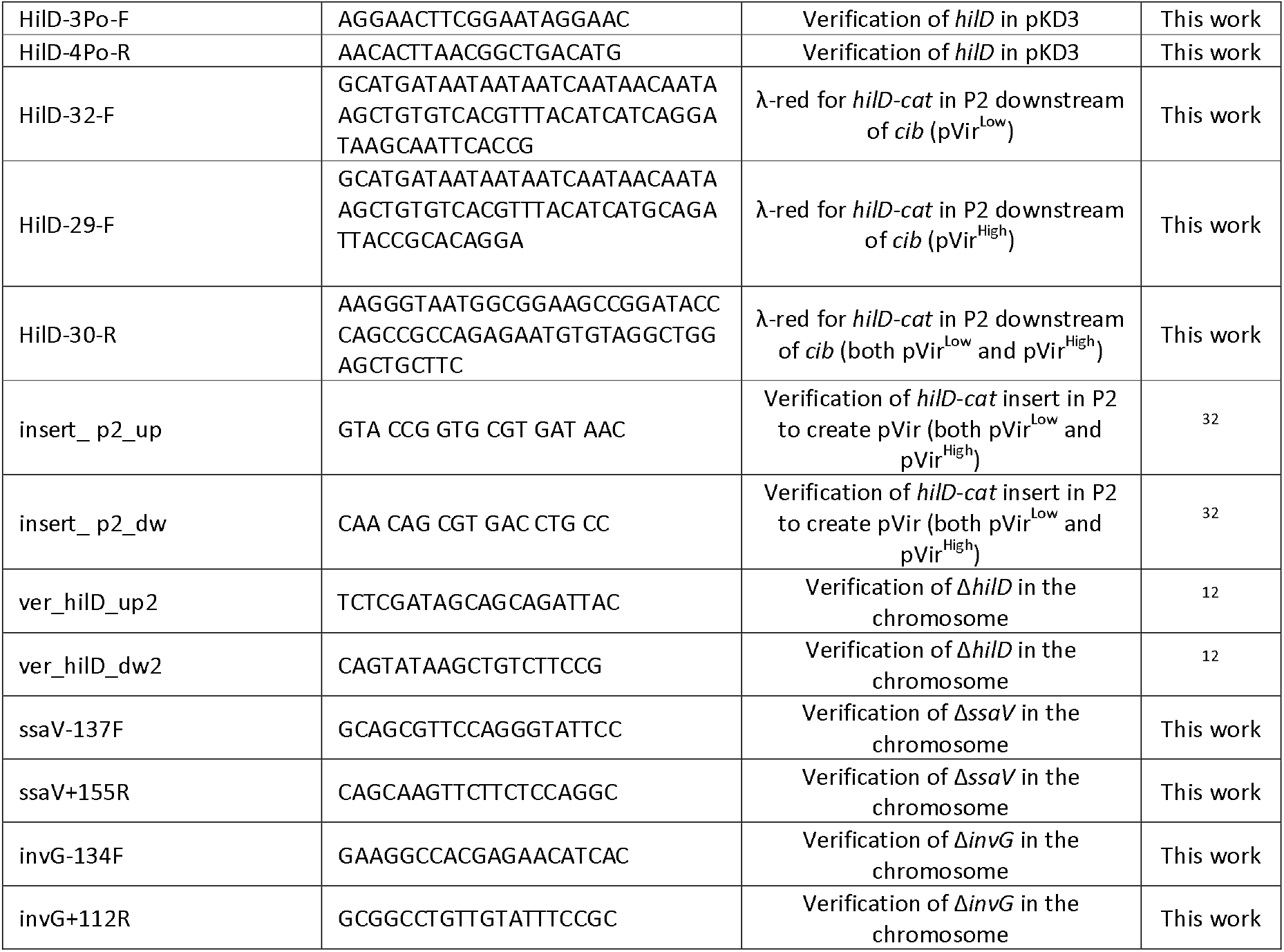
Primers used in this study.

### *In vitro* growth and *ttss-1* expression

Subcultures were grown in LB with appropriate antibiotics for 6 hours and subsequently diluted 200 times in 200 µl of media distributed in 96 well black side microplates (Costar). The lid-closed microplates were incubated at 37°C with fast and continuous shaking in a microplate reader (Synergy H4, BioTek Instruments). Optical density at 600 nm and GFP fluorescence (491 nm excitation; 512 nm emission) were measured every 10 minutes for 14 h. OD and fluorescence values were corrected for the baseline value measured for sterile broth.

### Infection experiments

All mouse experiment protocols are derived from the streptomycin pretreated mouse model described in^37^. We used ampicillin rather than streptomycin since *S*.Tm 14028S is not naturally resistant to streptomycin. Ampicillin resistance is conferred by pM975 contained in all strains used *in vivo*. All experiments were performed in 8-12 week old specified opportunistic pathogen free (SOPF) C57BL/6 mice, which were given 20 mg of ampicillin by oral gavage to allow robust colonization of *S*.Tm. This ampicillin pretreatment model has been used previously to measure HGT in the gut^30-32^. All infection experiments were approved by the responsible authorities (Tierversuchskommission, Kantonales Veterinäramt Zürich, licenses 193/2016 and 158/2019). Sample size was not predetermined and mice were randomly assigned to treatment group.

#### Single infections of donors or recipients

Overnight cultures grown at 37°C in LB with the appropriate antibiotics were diluted 1:20 and subcultured for 4 hours in LB without antibiotics. Cells were centrifuged and resuspended in PBS before being diluted. Ampicillin pretreated mice were orally gavaged with ∼5×10^7^ CFU. Fecal samples were collected daily, homogenized in PBS with a steel ball at 25 Hz for 1 minute, and bacterial populations were enumerated on selective MacConkey agar. Lipocalin-2 ELISA (R&D Systems kit; protocol according to manufacturer) was performed on feces to determine the inflammatory state of the gut. At day 4 post infection, mice were euthanized.

#### Plasmid transfer experiments

Donor and recipient strains (14028S derivatives; *ssaV* mutants) were grown overnight in LB with the appropriate antibiotics at 37°C and subsequently diluted 1:20 and subcultured for 4 hours in LB without antibiotics, washed in PBS, and diluted. Ampicillin pretreated mice were orally gavaged sequentially with ∼10^2^ CFU of donors followed by ∼10^4^ CFU of recipients. Feces were collected when needed, homogenized in PBS, diluted, and bacterial populations were enumerated on MacConkey agar containing the appropriate antibiotics (donors = Cm; recipients = Kan; transconjugants Cm+Kan; total population = populations of donors + recipients). Replica plating was used if the CFUs on the Cm+Kan plates approached those on the Cm or Kan plates to determine an exact ratio of plasmid transfer, and the donor population size. At day 10 post infection, mice were euthanized. Lipocalin-2 ELISA was performed on feces to determine the inflammatory state of the gut. When needed, transconjugants on the Cm+Kan plates were kept at 4°C until analysis by colony blot.

#### Competitions involving mobile versus non-mobile pVir

Cooperator (donor) and cheater (recipient) strains (14028S derivatives; *ssaV* mutants) were grown overnight in LB with the appropriate antibiotics at 37°C and subsequently diluted 1:20 for a 4 hour subculture in LB without antibiotics. Cells were resuspended in PBS and diluted. Ampicillin pretreated mice were sequentially orally gavaged with ∼10^2^ CFU of cooperators (pVir^Low^) immediately followed by ∼10^2^ CFU of cheaters. Feces were collected when needed, homogenized, diluted, and bacterial populations were enumerated on MacConkey agar containing the appropriate antibiotics as for the plasmid transfer experiments. At day 4 or 10 post infection, mice were euthanized. Lipocalin-2 ELISA was performed on feces to determine the inflammatory state of the gut. The colonies on the Cm plates from day 10 fecal plates were kept at 4°C until analysis by colony blot.

#### Transmission experiments

Feces from mice given donor and recipient strains were collected on day 2 and day 10 p.i., resuspended in PBS, briefly centrifuged, and 100 µl of the suspension was given to ampicillin pretreated mice. These experiments occurred in parallel to the plasmid transfer experiments to ensure fresh fecal populations were transmitted into new mice. Bacterial populations and the state of inflammation were measured as for the plasmid transfer experiments. Mice were euthanized at day 4 post transmission.

#### Evolved transconjugant infections

Single clones from plasmid transfer experiments were isolated on day 7 or day 10, and stored in 20% LB+glycerol at -80°C. Isolates were grown in LB containing the appropriate antibiotics (Cm, Kan, Amp) overnight at 37°C and subsequently diluted 1:20 and subcultured for 4 hours in LB without antibiotics. Of note, loss of pM975 was observed for some clones (based on loss of ampicillin resistance), and could therefore not be used for infection. Subcultured cells were centrifuged, resuspended in PBS, and ∼5×10^7^ CFU were given to ampicillin pretreated mice by oral gavage. The shedding population was enumerated on MacConkey supplemented with chloramphenicol after suspension in PBS followed by dilution. On day 4 post infection, mice were euthanized and fecal samples were additionally enumerated on MacConkey supplemented with kanamycin, to ensure that plasmid loss did not contribute to the detected shedding population. The kanamycin resistant colonies (Kan^R^ is encoded on the chromosome) were replica plated onto MacConkey supplemented with chloramphenicol to confirm that no pVir plasmid loss occurred. The MacConkey chloramphenicol plates were stored at 4°C until analysis by colony blot. LCN2 ELISA was used to determine the inflammatory state of the mice over time.

### Colony blots

To assess *ttss-1* expression at the clonal level (to determine the proportion of cooperators), a colony Western blot was performed. SipC was used as a proxy for *ttss-1* expression, since SipC is regulated by HilD. We have previously established this protocol to assess heterogeneously expressed phenotypes such as *ttss-1* in *S*.Tm^12,24^, since single-cell approaches would not differentiate cheaters from the phenotypically OFF subpopulation^24^. For a detailed protocol and an overview of applications, see^24^. Briefly, colonies on MacConkey agar were replica transferred to nitrocellulose membranes and placed face-up on LB agar without antibiotics and allowed to grow overnight. The original MacConkey plates are also allowed to re-grow and then stored at 4°C. Colonies were lysed and cellular material was hybridized to the membrane by passing the membranes over a series of Whatman filter papers soaked with buffers: 10 minutes on 10% SDS, 10 minutes on denaturation solution (0.5 M NaOH, 1.5 M NaCl), twice for 5 minutes on neutralization solution (1.5 M NaCl, 0.5 M Tris-HCl, pH 7.4), and 15 minutes on 2× SSC (3 M NaCl, 0.3 M sodium citrate, pH 7). Membranes were washed twice with TBS (10 mM Tris-HCl, 150 mM NaCl, pH 7.4) and excess cellular debris was gently removed by scraping the surface with a folded Whatman paper. Membranes were blocked with TBS containing 3% BSA for 1 hour at room temperature and then incubated with 5 ml of TBS with 3% BSA containing a 1:4000 dilution of anti-SipC rabbit antibody provided by Virotech Diagnostics GmbH (reference number: VT110712) overnight in a moist chamber at 4°C on a rocking platform. Washing once with TBS-T (20 mM Tris-HCl, 500 mM NaCl, 0.05% Tween 20, 0.2% Triton X-100, pH 7.5) and twice with TBS removed non-specific binding. Secondary antibodies (1:2500 dilution of goat anti-rabbit IgG conjugated to HRP; Sigma; catalogue number A0545-1ML) were then added to membrane in TBS with 3% BSA and incubated at room temperature on a rocking platform for 2-4 hours. Three more washing steps with TBS were performed before resolving the staining with 5 ml of substrate per membrane: a 30 mg tablet of 4-chloro-1-naphthol (Sigma) dissolved in 10 ml of methanol, mixed with H^2^O^2^ (0.06% w/v) in 50 ml of TBS. The reaction is stopped with water after the desired intensity is observed.

Clones of interest can be identified by changes in SipC abundance. Desired isolates were matched to the original MacConkey plate and inoculated in LB containing chloramphenicol and kanamycin. Isolates were then stored in 20% LB+glycerol at -80°C until whole-genome bacterial sequencing was performed, or evolved clones were used for infection.

### Whole-genome bacterial sequencing

Strains stored in 20% LB+glycerol at -80°C were inoculated in LB with the appropriate antibiotics. Genomic DNA was extracted from 1 ml of overnight culture using a QIAamp DNA Mini Kit (Qiagen). Illumina MiSeq sequencing operated by the Functional Genomics Centre Zurich and Novogene (Cambridge) was performed to generate 150bp paired end reads with at least 50× coverage across the genome. Bioinformatic analysis was performed using CLC Genomics Workbench 11.0. Reads were mapped to the 14028S chromosome reference (NCBI accession NC_016856.1) and the pVir plasmids (the SL1344 P2 plasmid (NCBI accession NC_017718.1) was modified by inserting the cloned *hilD-cat* regions to create pVir^Low^ and pVir^High^ reference sequences). Basic variant detection was performed to detect variants that occurred in a minimum of 70% of reads. Variants were excluded if they occurred in non-specific regions determined by read mapping in CLC (e.g. where reads could map equally well to another location in the genome). Small insertions or deletions (Indels) were also detected using software in CLC Genomics Workbench. This is summarized in **Tables S1-4**.

### Statistical analysis

Statistical tests on experimental data were performed using GraphPad Prism 8 for Windows.

## Acknowledgements

We would like to thank members of the Hardt and Diard labs for valuable discussions, as well as the staff at RCHCI and EPIC animal facilities. We thank Stuart West, Kevin Foster and Ashleigh Griffin for helpful feedback and comments on this manuscript. We would also like to acknowledge the staff at Functional Genomics Centre Zurich and Novogene for whole-genome bacterial sequencing. We acknowledge grant funding from the Swiss National Science Foundation (NRP 72 407240_167121, 310030B_173338 and 310030_192567), the Gebert Rüf Foundation (GRS-060/18) and the Monique Dornonville de la Cour Foundation to WDH. MD is funded by an SNF professorship grant (PP00PP_176954) and a BRCCH multi-investigator grant, JSH is funded by NRP72 grant 407240- 167121 from the Swiss National Science Foundation, and EB received a Boehringer Ingelheim Fonds PhD fellowship. The authors declare no conflict of interest.

## Contributions

EB, WDH, and MD conceived the project and designed the experiments. EB, EG, YS, and AR carried out the experiments and analyzed data. EB, WDH, and MD wrote the manuscript. EG and JSH provided valuable input on experimental design, theoretical background, and the manuscript. All authors read, commented on, and approved this manuscript.

## Supplementary information

### Supplementary discussion: Why does inflammation cause only mild *S*.Tm blooms as detected by shedding in Fig. 1?

In our experimental system, we address if HGT can facilitate the emergence of cooperative virulence using *S*.Tm as a model system. *S*.Tm is an excellent candidate to monitor the emergence of cooperative virulence since tractable models for within-host evolution of *S*.Tm (e.g. plasmid transfer and evolution of virulence) have been established that integrate important parameters of the host- pathogen interaction^4,8,37,41,61^. The antibiotic pretreatment model allows robust colonization of *S*.Tm in mice, leading to inflammation that acts as a public good to suppress the microbiota and create a favourable niche for Enterobacteriaceae, such as *S*.Tm, to thrive^8,37^. In the absence of overt inflammation, the microbiota can re-grow and exclude *S*.Tm from the gut^38^.

Our experimental approach uses this antibiotic pretreatment model to allow efficient gut luminal colonization of *S*.Tm, a pre-requisite for efficient plasmid exchange. In the experiments in **Fig. 1**, one day after ampicillin pretreatment, two avirulent *S*.Tm strains are introduced into the gut and allowed to grow and exchange plasmids. This experimental setup is unique in the sense that for the first time using this mouse model, we introduce virulence only after plasmid exchange, meaning that virulence can emerge from rare. Virulent clones are initially very low in abundance and only expand to sufficient density to trigger inflammation (∼10^8^ CFU/g feces^9^) at day 2 post infection (**Fig. 2B**), mediated by plasmid spread. This means that there is a delay of approximately 1-2 days in triggering inflammation, compared to the conventional antibiotic pretreatment model. Microbiota re-growth then likely competes against actively invading *S*.Tm that have recently become virulent, creating a delicate balance that may ultimately prevent overt inflammation. Indeed, the inflammation measured by LCN2 ELISA ranges from 10^1^ - 10^3^ ng LCN2 / g feces (median ∼10^2^ ng / g feces at day 3 post infection; **Fig. 1C**), which is strikingly lower than the inflammation triggered in a scenario where virulent clones are present immediately after introduction in the gut (e.g. **Fig. 3D; Fig. 4B**,**E; Fig. 5A**; ∼10^3^ - 10^4^ ng LCN2 / g feces at day 1 post infection). The lack of immediate overt inflammation in **Fig. 1C** can explain the absence of clear pathogen blooms (**Fig. 1D**) typically associated with inflammation^8^ (i.e., the microbiota starts to re-grow and cannot be suppressed by mild inflammation). This is in contrast to when shedding is maintained in mice where overt inflammation is detected (e.g. **Fig. 3E; Fig. 4C**,**F; Fig. 5B**).

What does this mean for the emergence of virulence by HGT? The intuition for the prevalence of virulence factors in pathogens such as *S*.Tm is that they provide a selective advantage, since HGT-mediated maintenance alone is likely too unstable to be the sole driver for the evolution of virulence factors **(Fig. 2A; Fig. 3G; Fig. 4D**,**G; Fig. 5C;**^15,18^). However, it is also unlikely that HGT of a virulence-conferring allele permeates the pathogen population immediately to lead to a discernable fitness advantage in the initial host (e.g. pVir spreading to 100% of the population immediately to evoke inflammation-mediated blooms). Therefore, the selective advantage must be elsewhere. Here, we show that pathogen cells that harbour newly acquired virulence determinants can lead to inflammation after transmission, even if inflammation was not overtly triggered in the initial host (e.g. compare inflammation and shedding in **Fig. 1** to **Fig. 3**. Therefore, the lack of pathogen blooms in the first host can be compensated with the inflammation triggered in subsequent hosts after transmission.

**Supplementary Figure S1.**
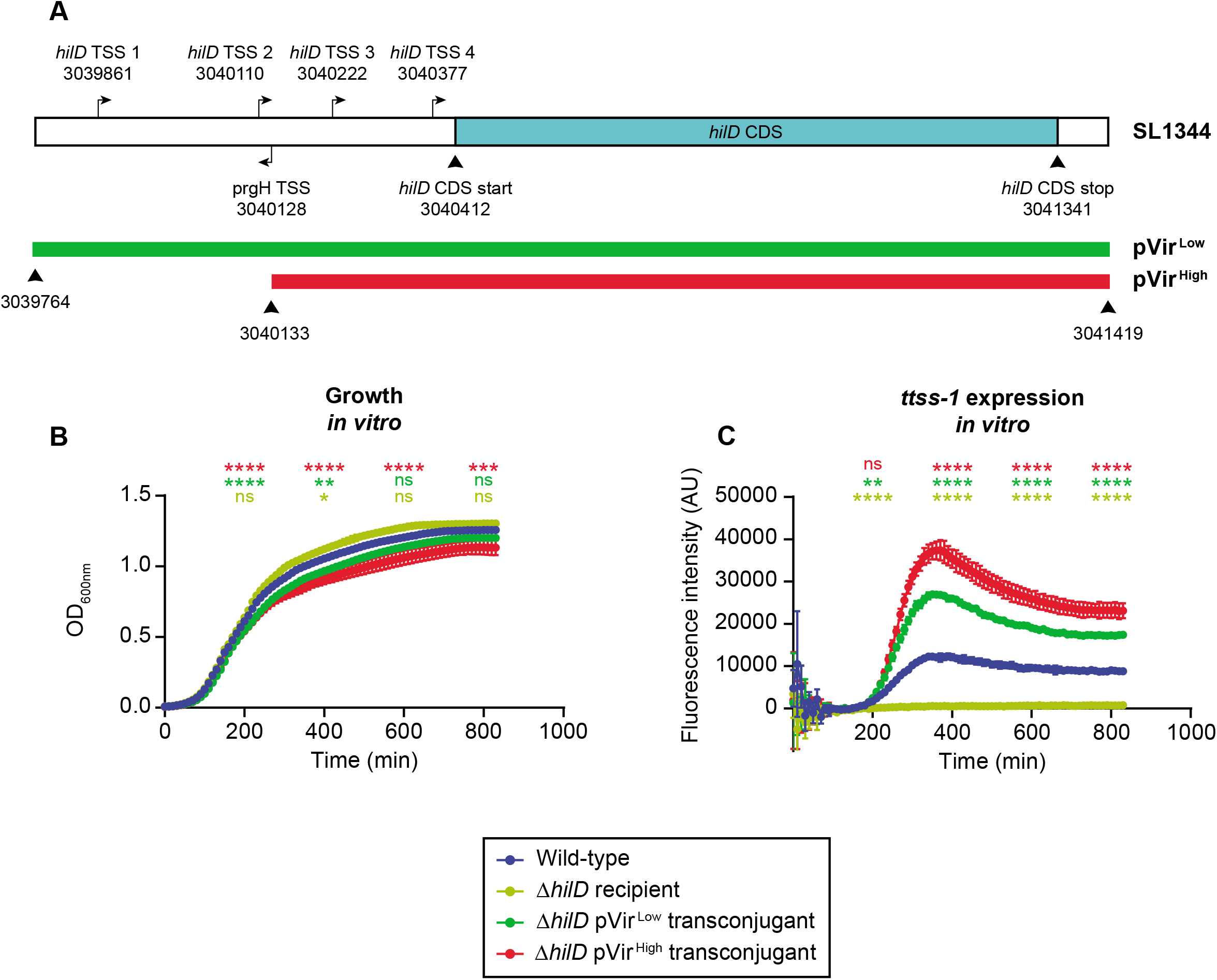
Genetic locus of *hilD* cloned into pVir and characterization of *ttss-1* expression and cost *in vitro*. **A**) The *hilD* coding sequence (CDS) of *S*.Tm SL1344 (NCBI Accession: FQ312003.1), the template for cloning the two pVir constructs, is shown along with the four transcriptional start sites (TSS) of *hilD* identified in^36^ (indicated with arrows; the *prgH* TSS is also indicated). The position in the SL1344 genome is indicated numerically. The regions cloned into pVir^Low^ or pVir^High^ are indicated with a green or red bar, respectively. Positions along the 1655 bp genetic locus shown are to scale. **B-C)** *In vitro* analysis of pVir^Low^ (green) and pVir^High^ (red) transconjugants compared to a wild-type (blue) and a Δ*hilD* mutant (yellow). Strains bearing pM972 (*P*sicA*-gfp*; *ttss-1* expression reporter) were used to correlate growth (**panel B**) and *ttss-1* expression (**panel C**). Data is shown as the mean with standard deviation of 2 independent clones for pVir^Low^, 6 independent clones for pVir^High^, 1 clone for wild-type and 1 clone for the Δ*hilD* mutant (3 technical replicates for each clone). A one-way ANOVA with Dunnett’s multiple comparisons test is performed at 200, 400, 600, and 800 min post inoculation, comparing each group to the wild-type control (indicated with asterisks corresponding to the colour of the group; p>0.05 (ns), p<0.05 (*), p<0.01 (**), p<0.001 (***), p<0.0001 (****)). All strains lacked functional *ttss-2* genes (*ssaV* mutation).

**Supplementary Figure S2.**
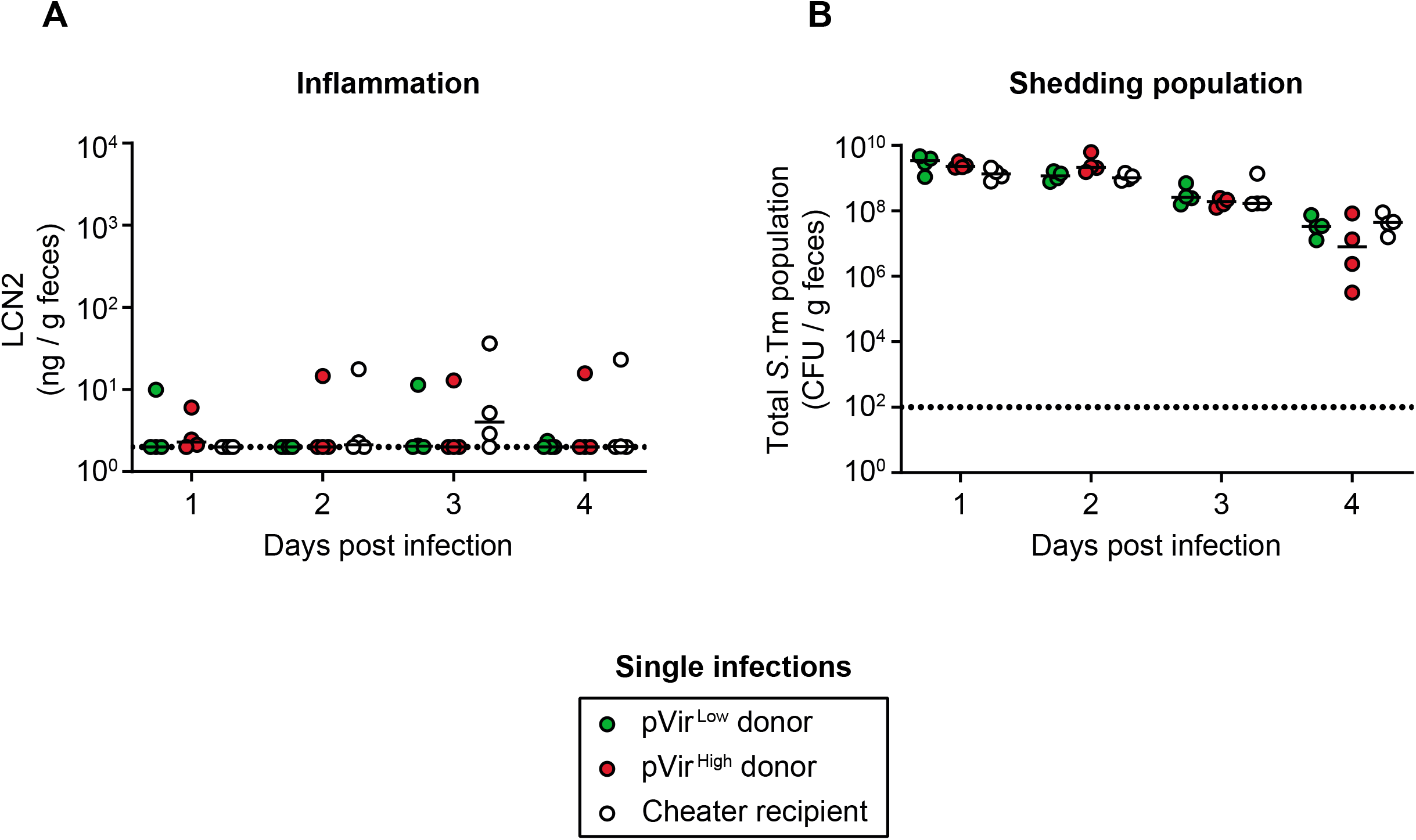
The pVir^Low^ and pVir^High^ donors and the recipient cannot trigger inflammation during single-infections. Ampicillin pretreated mice (n=4 per group) were orally infected, either with ∼5×10^7^ CFU of pVir^Low^ (green) donors (14028S Δ*invG* Δ*hilD* Δ*ssaV* pVir), pVir^High^ (red) donors (14028S Δ*invG* Δ*hilD* Δ*ssaV* pVir), or with recipients (14028S Δ*hilD* Δ*ssaV*; white). All data points are shown and medians are indicated with lines for **A)** Inflammation detected by a LCN2 ELISA and **B)** the total *S*.Tm population quantified by selective plating. Dotted lines indicate the detection limit.

**Supplementary Figure S3.**
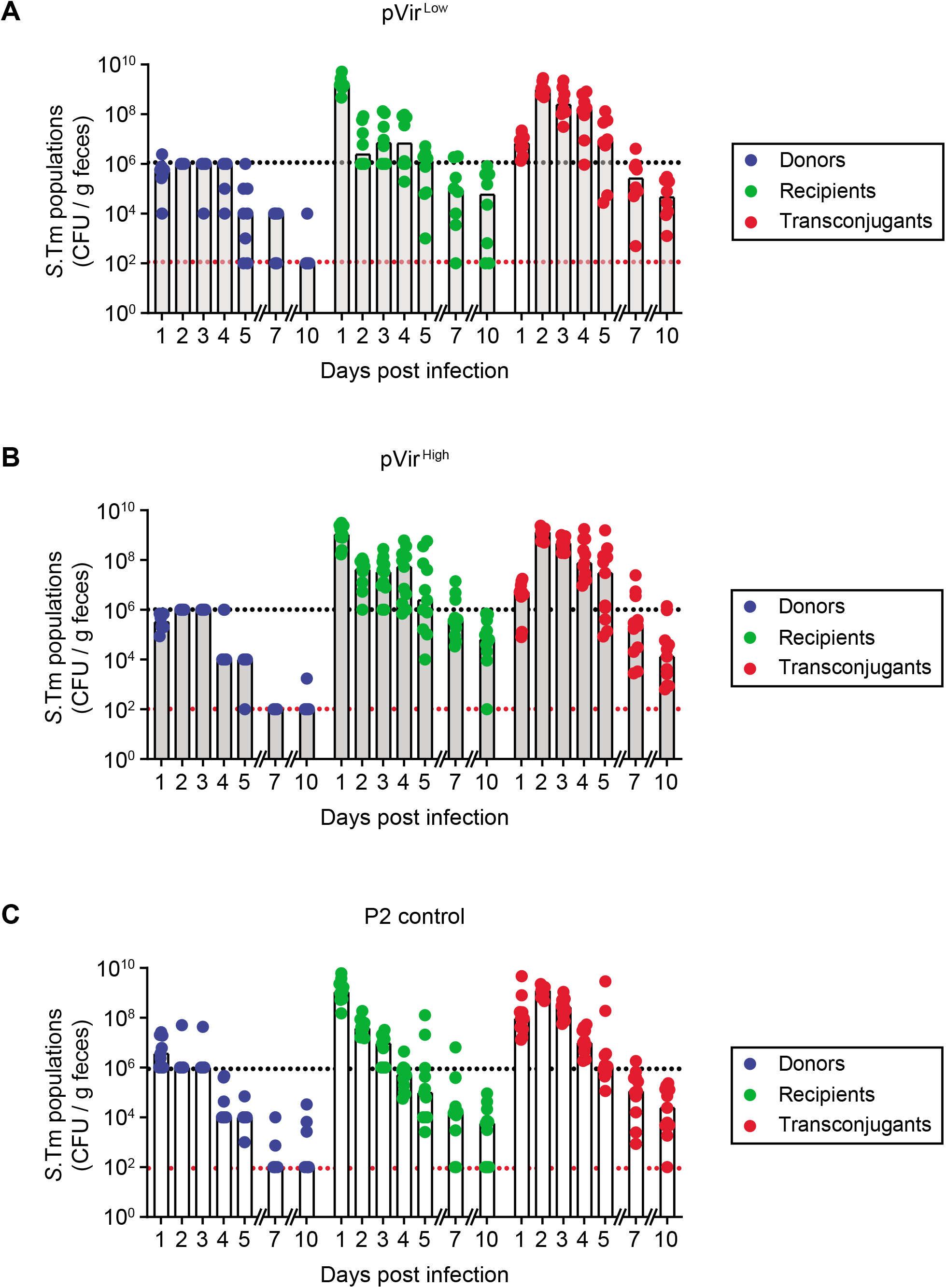
Fecal bacterial population sizes in samples from experiments in figure 1B-C. Fecal loads of donors (blue; Cm^R^), recipients (green; Kan^R^), and transconjugants (red; Cm^R^, Kan^R^) determined by selective plating. Replica plating was used to determine exact ratios of transconjugants compared to either donors or recipients. The black dotted line indicates the conservative detection limit for donors and recipients (depending on the dilution used for replica plating, values can appear below this line), and the red dotted line indicates the detection limit for transconjugants. Each data point is represented and bars indicate the median. **A)** Mice infected with *S*.Tm donors carrying pVir^Low^. **B)** Mice infected with *S*.Tm donors carrying pVir^High^. **C)** Mice infected with *S*.Tm donors with P2.

**Supplementary Figure S4.**
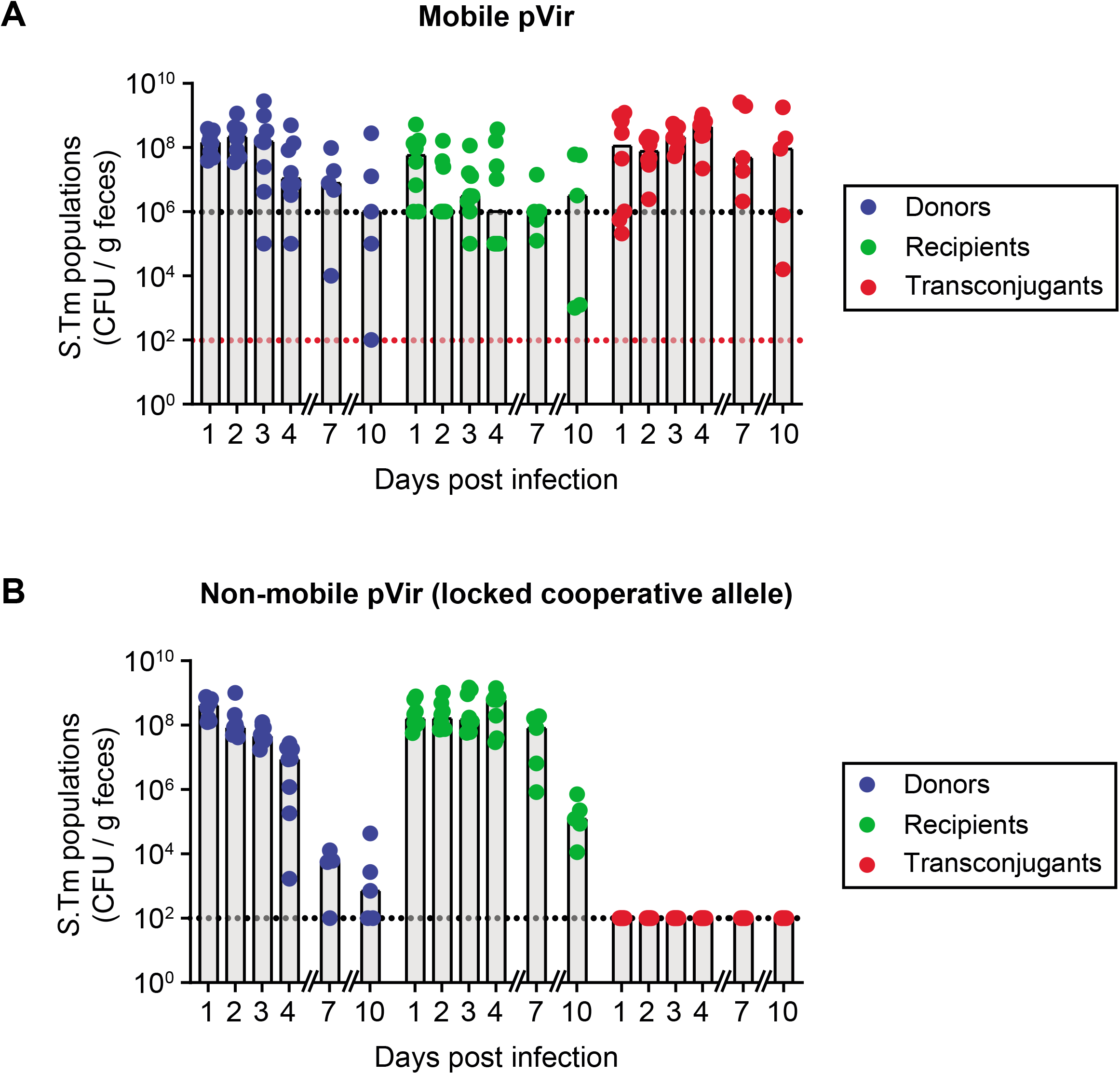
Fecal bacterial population sizes in samples from experiments in figure 3B-G. Fecal loads of donors (blue; Cm^R^), recipients (green; Kan^R^), and transconjugants (red; Cm^R^, Kan^R^) determined by selective plating. Replica plating was used to determine exact ratios of transconjugants compared to either donors or recipients. Each data point is represented and bars indicate the median. **A)** Mice infected with *S*.Tm donors and recipients in the mobile scenario. The black dotted line indicates the conservative detection limit for donors and recipients (depending on the dilution used for replica plating, values can appear below this line), and the red dotted line indicates the detection limit for transconjugants. **B)** Mice infected with *S*.Tm donors and recipients in the non-mobile scenario. The black dotted line indicates the detection limit.

**Supplementary Figure S5.**
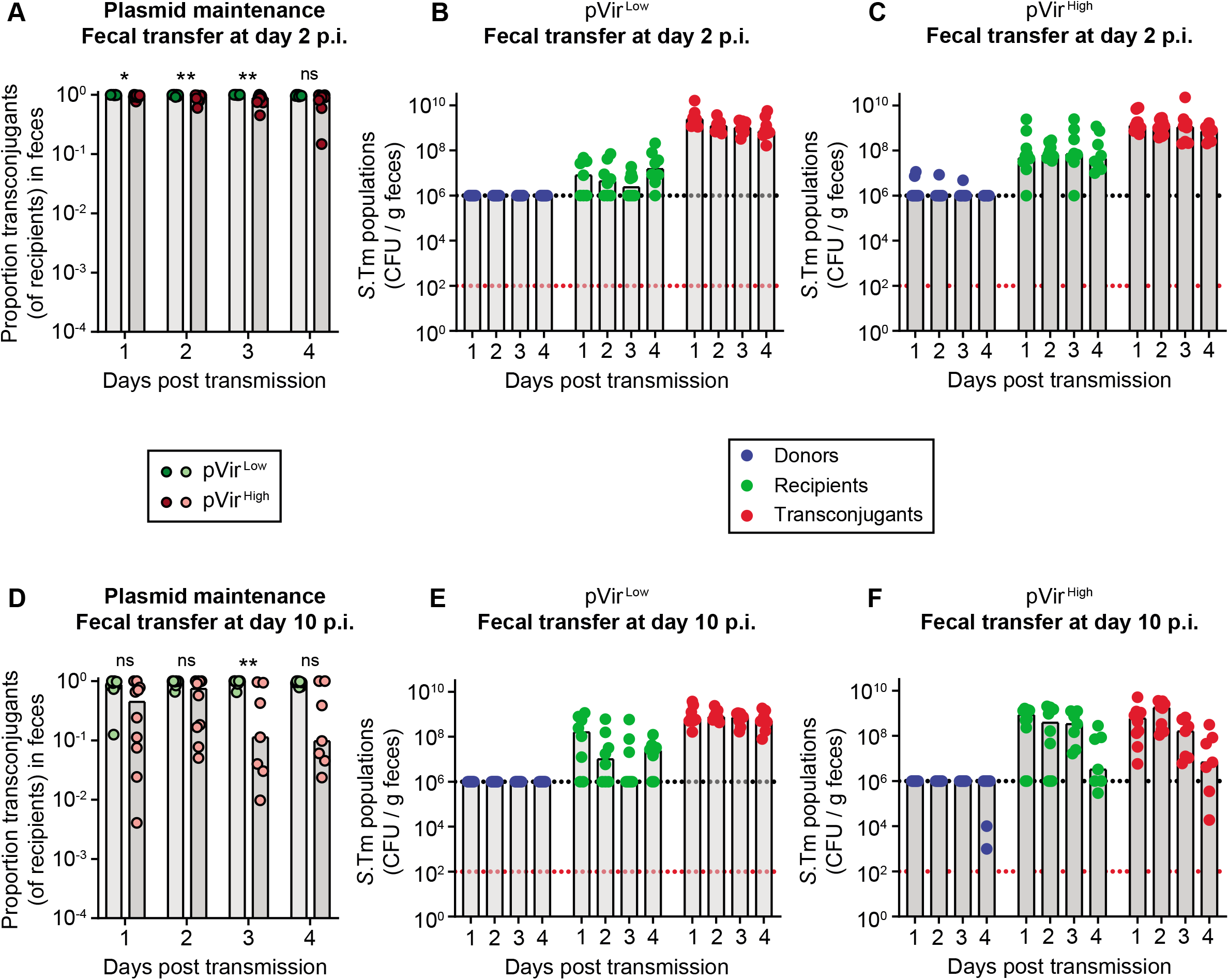
Plasmid maintenance and fecal bacterial population sizes of mice in Figure 3B-G. **A-C)** Mice transmitted with fecal suspensions from day 2 post infection. **D-F)** Mice transmitted with fecal suspensions from day 10 post infection. **A**,**D)** Plasmid transfer was measured by selective plating: donors Cm^R^, recipients Kan^R^, and transconjugants both Cm^R^ and Kan^R^. The proportion of transconjugants is calculated by dividing the transconjugant population by the sum of recipients and transconjugants. All data points are shown and medians are indicated by bars. Replica plating was used to determine exact ratio of transconjugants compared to recipients. Mice given fecal resuspensions with *S*.Tm harbouring pVir^Low^ (green; dark shade for day 2 transmission; light shade for day 10 transmission) are compared to pVir^High^ (red; dark shade for day 2 transmission; light shade for day 10 transmission) using a two-tailed Mann-Whitney U test (p>0.05 (ns), p<0.05 (*), p<0.01 (**), p<0.001 (***), p<0.0001 (****)). **B**,**C**,**E**,**F)** Fecal loads of donors (blue), recipients (green), and transconjugants (red) determined by selective plating. The black dotted line indicates the conservative detection limit for donors and recipients (depending on the dilution used for replica plating, values can appear below this line), and the red dotted line indicates the detection limit for transconjugants. Each data point is represented and bars indicate the median. **B**,**E)** Mice transmitted with fecal suspensions containing *S*.Tm with pVir^Low^. **C**,**F)** Mice transmitted with fecal suspensions containing *S*.Tm with pVir^High^.

**Supplementary Figure S6.**
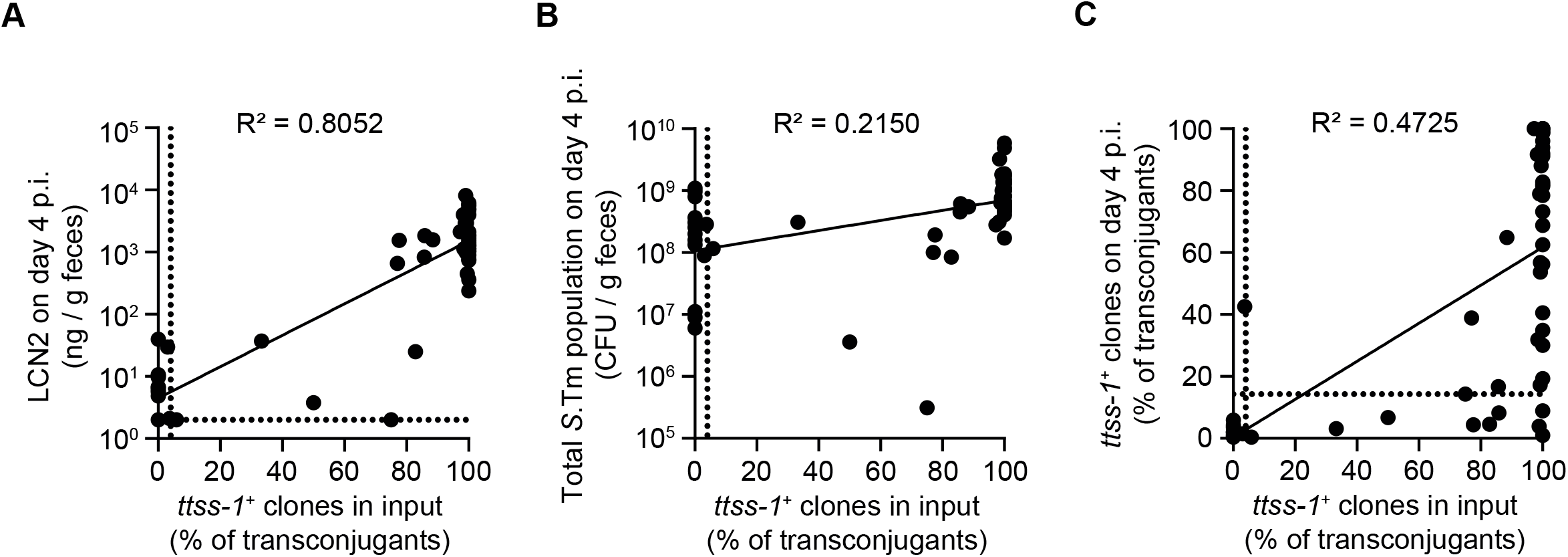
The proportion of cooperative clones transmitted to mice can predict disease, shedding, and the cheating dynamics. The proportion of cooperative clones (*ttss-1*^*+*^ clones) given to mice (x-axis; determined from **Fig. 2A**; assumed to be 100% for a *ttss-1*^*+*^ evolved clone and 0% for a *ttss-1*^*-*^ evolved clone) is plotted against the resulting inflammation (*panel A*), shedding population (*panel B*), and proportion of cooperative clones (*panel C*) at day 4 post infection in mice from **Fig. 3** and **Fig. 4** (y-axis). A linear regression was performed (y-axis log transformed in **panels A**,**B**) and was significantly non-zero as determined with an F test (p<0.0001 for **panels A**,**C**; p=0.0002 for **panel B**). The line of best fit and the goodness of fit (R^2^) is shown on the graphs. Dotted lines indicated the detection limits. For detection limits based on colony blots, the conservative detection limit is shown, which is dependent on the number of colonies on the plate (values can therefore appear below the detection limit).

**Supplementary Figure S7.**
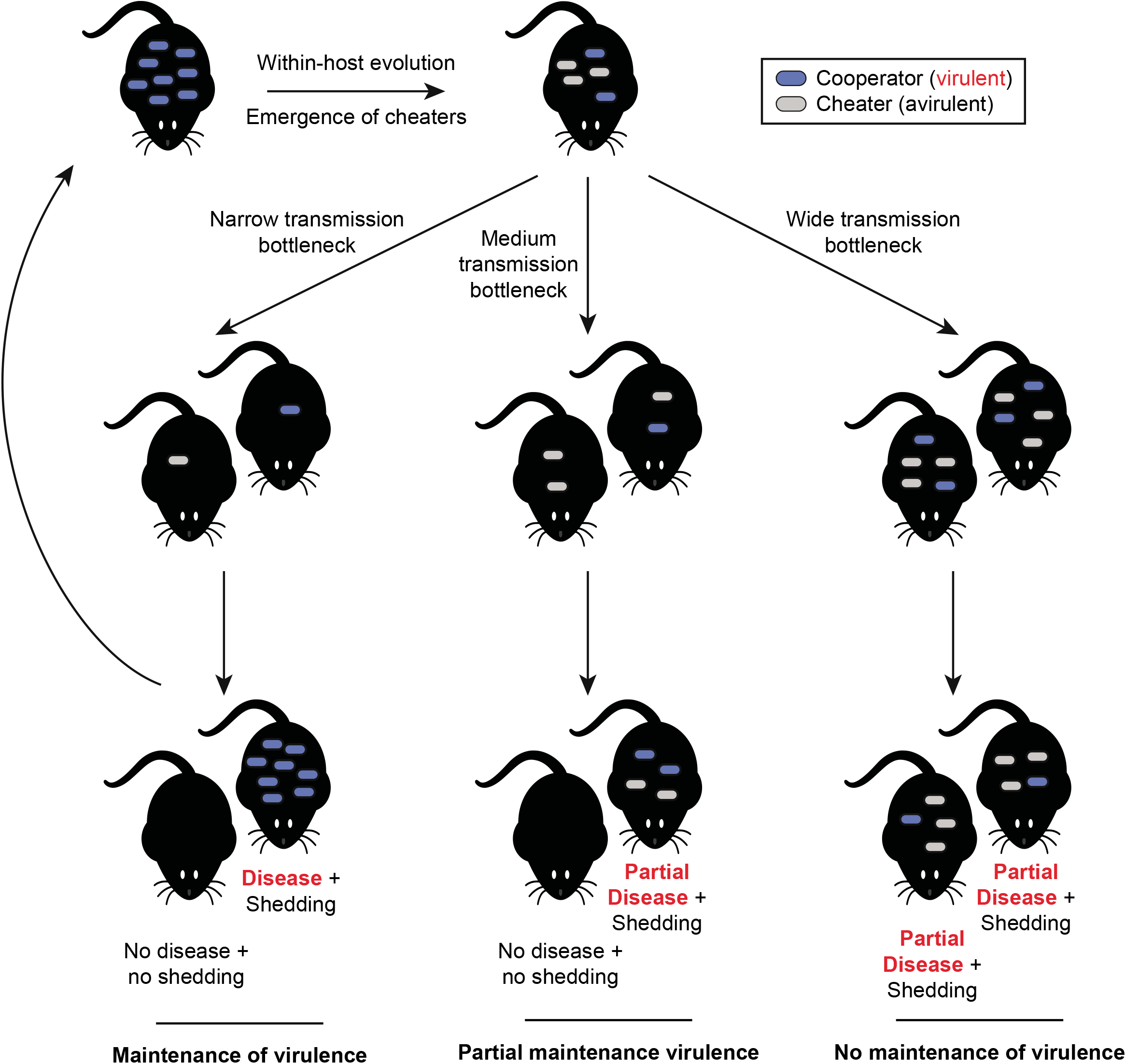
Proposed model for how transmission bottlenecks can select for cooperative virulence in *S*.Tm. Cheaters can emerge and expand in the population, leading to a destabilization of cooperation. Establishment in the next host is dependent on the proportion of cooperators. Wide or medium transmission bottlenecks (such as in **Fig. 4**) allow for cheaters to free-ride off of the inflammation triggered by cooperating clones. Since they will also outcompete cooperating clones, the shedding is determined by the amount of cooperating clones initially. In cases where only cheating clones are transmitted, shedding cannot be achieved because inflammation is not triggered. Only in cases of narrow bottlenecks, which are more likely in nature, cooperating clones can be transmitted in the absence of cheating clones, leading to inflammation-associated blooms and shedding. This allows evolution of virulence.

**Table S1. Summary of SNPs or indels in pVir**^**Low**^ **in evolved transconjugants**. For all clones, the SipC phenotype from colony blot (as a proxy for *ttss-1* expression; + indicates positive and blue fill labels clones of that phenotype, - indicates negative and yellow fill labels clones of that phenotype) is shown and the mutations are summarized. If blank, the region of the genome contains no mutations. The *hilD* genotype is highlighted in either blue or yellow fill according to the SipC phenotype to allow for clear correlation.

**Table S2. Summary of SNPs or indels in pVir**^**High**^ **in evolved transconjugants**. For all clones, the SipC phenotype from colony blot (as a proxy for *ttss-1* expression; + indicates positive and blue fill labels clones of that phenotype, - indicates negative and yellow fill labels clones of that phenotype) is shown and the mutations are summarized. If blank, the region of the genome contains no mutations. The *hilD* genotype is highlighted in either blue or yellow fill according to the SipC phenotype to allow for clear correlation.

**Table S3. Summary of SNPs or indels in coding sequences of the chromosome of evolved transconjugants with pVir**^**Low**^. For all clones, the SipC phenotype from colony blot (as a proxy for *ttss- 1* expression; + indicates positive and blue fill labels clones of that phenotype, - indicates negative and yellow fill labels clones of that phenotype) is shown and the mutations are summarized. For variants that affect single nucleotides, the position in the 14028S reference chromosome (NCBI accession NC_016856.1) is indicated along with the allele change. For targeted deletions introduced by allelic replacement or P22 transduction, the “Δ” symbol is used. Variants are only shown if they occurred in >70% of reads. Mutations are excluded if they also occurred in our ancestral lab strain of 14028S. If blank, the region of the genome contains no mutations.

**Table S4. Summary of SNPs or indels in coding sequences of the chromosome of evolved transconjugants with pVir**^**High**^. For all clones, the SipC phenotype from colony blot (as a proxy for *ttss- 1* expression; + indicates positive and blue fill labels clones of that phenotype, - indicates negative and yellow fill labels clones of that phenotype) is shown and the mutations are summarized. For variants that affect single nucleotides, the position in the 14028S reference chromosome (NCBI accession NC_016856.1) is indicated along with the allele change. For targeted deletions introduced by allelic replacement or P22 transduction, the “Δ” symbol is used. Variants are only shown if they occurred in >70% of reads. Mutations are excluded if they also occurred in our ancestral lab strain of 14028S. If blank, the region of the genome contains no mutations.

