## Supplemental Tables for "Cooperative virulence can emerge via horizontal gene transfer but is stabilized by transmission"

**Table S1. Summary of SNPs or indels in pVir^Low^ in evolved transconjugants.**

|  | Clone | Ancestral donor pVir^Low^ | Z2296 | Z2298 | Z2299 | Z2301 | Z2302 | Z2304 | Z2305 | Z2306 | Z2308 | Z2309 | Z2310 |
| --- | --- | --- | --- | --- | --- | --- | --- | --- | --- | --- | --- | --- | --- |
|  | **pVir type** | Low | Low | Low | Low | Low | Low | Low | Low | Low | Low | Low | Low |
|  | **Day isolated** | N/A | 7 | 7 | 7 | 7 | 7 | 7 | 10 | 10 | 10 | 7 | 7 |
|  | **SipC phenotype** | - | + | - | + | - | + | - | - | + | + | - | + |
|  | **Mouse** | N/A | AL876 | AL876 | AL877 | AL877 | AL878 | AL878 | AM128 | AM128 | AM129 | AM130 | AM130 |
| pVir mutations | **Annotated function** |  |  |  |  |  |  |  |  |  |  |  |  |
| *hilD* | transcriptional regulator HilD |  |  | 285 bp Indel |  | 134 bp Indel |  | 409 bp Indel | 100 bp Indel |  |  | 365 bp Indel  G>A (pos 584 in CDS) |  |

**Table S2. Summary of SNPs or indels in pVir^High^ in evolved transconjugants.**

|  | Clone | Ancestral donor pVir^High^ | Z2238 | Z2239 | Z2242 | Z2243 | Z2244 | Z2245 | Z2246 | Z2247 | Z2252 | Z2253 | Z2254 | Z2255 | Z2311 | Z2312 |
| --- | --- | --- | --- | --- | --- | --- | --- | --- | --- | --- | --- | --- | --- | --- | --- | --- |
|  | **pVir type** | High | High | High | High | High | High | High | High | High | High | High | High | High | High | High |
|  | **Day isolated** | N/A | 10 | 10 | 10 | 10 | 10 | 10 | 10 | 10 | 10 | 10 | 10 | 10 | 10 | 10 |
|  | **SipC phenotype** | - | + | - | + | - | + | - | + | - | - | + | - | - | - | + |
|  | **Mouse** | N/A | AG58 | AG60 | AG300 | AG300 | AG301 | AG301 | AG302 | AG302 | AG590 | AG591 | AG591 | AG592 | AM132 | AM132 |
| pVir mutations | **Annotated function** |  |  |  |  |  |  |  |  |  |  |  |  |  |  |  |
| *hilD* | transcriptional regulator HilD |  |  | 5 bp Indel |  | 63 bp Indel |  | 63 bp Indel |  | G > A (pos 173 in CDS) |  |  |  | 133 bp Indel | 258 bp Indel |  |
| Upstream of *hilD* | transcriptional regulator HilD |  |  |  |  |  |  |  |  |  | 310 bp Indel |  | 326 bp Indel |  |  |  |
| SL1344_RS24670 | Transcription termination factor NusG |  |  |  | G > T (pos 415 in CDS) | G > T (pos 415 in CDS) | G > T (pos 415 in CDS) | G > T (pos 415 in CDS) | G > T (pos 415 in CDS) |  |  |  |  |  |  |  |

**Table S3. Summary of SNPs or indels in coding sequences of the chromosome of evolved transconjugants with pVir^Low^.**

|  | Clone | Ancestral donor pVir^Low^ | Ancestral recipient | Z2296 | Z2298 | Z2299 | Z2301 | Z2302 | Z2304 | Z2305 | Z2306 | Z2308 | Z2309 | Z2310 |
| --- | --- | --- | --- | --- | --- | --- | --- | --- | --- | --- | --- | --- | --- | --- |
|  | **pVir type** | Low | N/A | Low | Low | Low | Low | Low | Low | Low | Low | Low | Low | Low |
|  | **Day isolated** | N/A | N/A | 7 | 7 | 7 | 7 | 7 | 7 | 10 | 10 | 7 | 7 | 7 |
|  | **SipC phenotype** | **-** | **-** | + | - | + | - | + | - | - | + | + | - | + |
|  | **Mouse** | N/A | N/A | AL876 | AL876 | AL877 | AL877 | AL878 | AL878 | AM128 | AM128 | AM129 | AM130 | AM130 |
| Chromosomal mutations in CDS not present in the 14028 ancestor | **Annotated function** |  |  |  |  |  |  |  |  |  |  |  |  |  |
| *melR* | transcriptional regulator MelR |  |  |  | 4554843  G > A |  |  |  |  |  |  |  |  |  |
| *melB* | melbiose:sodium transporter MelB |  |  | 4558548  G > - |  | 4558548  G > - | 4558548  G > - | 4558548  G > - |  |  |  |  |  |  |
| *ssaV* | SPI-2 type III secretion system apparatus protein SsaV | Δ | Δ | Δ | Δ | Δ | Δ | Δ | Δ | Δ | Δ | Δ | Δ | Δ |
| *hilD* | transcriptional regulator HilD | Δ | Δ | Δ | Δ | Δ | Δ | Δ | Δ | Δ | Δ | Δ | Δ | Δ |
| *sipA* | SPI-1 type III secretion system effector SipA |  | 3046007  A > G | 3046007  A > G | 3046007  A > G | 3046007  A > G | 3046007  A > G | 3046007  A > G | 3046007  A > G | 3046007  A > G | 3046007  A > G | 3046007  A > G | 3046007  A > G | 3046007  A > G |
| *invG* | SPI-1 type III secretion system outer membrane ring protein InvG | Δ |  |  |  |  |  |  |  |  |  |  |  |  |

**Table S4. Summary of SNPs or indels in coding sequences of the chromosome of evolved transconjugants with pVir^High^.**

|  | Clone | Ancestral donor pVir^High^ | Ancestral recipient | Z2238 | Z2239 | Z2242 | Z2243 | Z2244 | Z2245 | Z2246 | Z2247 | Z2252 | Z2253 | Z2254 | Z2255 | Z2311 | Z2312 |
| --- | --- | --- | --- | --- | --- | --- | --- | --- | --- | --- | --- | --- | --- | --- | --- | --- | --- |
|  | **pVir type** | High | N/A | High | High | High | High | High | High | High | High | High | High | High | High | High | High |
|  | **Day isolated** | N/A | N/A | 10 | 10 | 10 | 10 | 10 | 10 | 10 | 10 | 10 | 10 | 10 | 10 | 10 | 10 |
|  | **SipC phenotype** | - | - | + | - | + | - | + | - | + | - | - | + | - | - | - | + |
|  | **Mouse** | N/A | N/A | AG58 | AG60 | AG300 | AG300 | AG301 | AG301 | AG302 | AG302 | AG590 | AG591 | AG591 | AG592 | AM132 | AM132 |
| Chromosomal mutations in CDS not present in the 14028 ancestor | **Annotated function** |  |  |  |  |  |  |  |  |  |  |  |  |  |  |  |  |
| *melR* | transcriptional regulator MelR |  |  |  |  |  |  |  |  |  |  | 4555269  T > G |  |  |  |  |  |
| *melB* | melbiose:sodium transporter MelB |  |  | 4558394  C > T |  |  | 4558406  T > G |  | 4558406  T > G |  |  |  | 4558403  G > A |  | 4558406  T > G |  | 4558156  T > A |
| *ssaV* | SPI-2 type III secretion system apparatus protein SsaV | Δ | Δ | Δ | Δ | Δ | Δ | Δ | Δ | Δ | Δ | Δ | Δ | Δ | Δ | Δ | Δ |
| *hilD* | transcriptional regulator HilD | Δ | Δ | Δ | Δ | Δ | Δ | Δ | Δ | Δ | Δ | Δ | Δ | Δ | Δ | Δ | Δ |
| *sipA* | SPI-1 type III secretion system effector SipA |  | 3046007  A > G | 3046007  A > G | 3046007  A > G | 3046007  A > G | 3046007  A > G | 3046007  A > G | 3046007  A > G | 3046007  A > G | 3046007  A > G | 3046007  A > G | 3046007  A > G | 3046007  A > G | 3046007  A > G | 3046007  A > G | 3046007  A > G |
| *invG* | SPI-1 type III secretion system outer membrane ring protein InvG | Δ |  |  |  |  |  |  |  |  |  |  |  |  |  |  |  |
| *tatD* | 3'-5' ssDNA/RNA exonuclease TatD |  |  |  |  |  |  |  |  |  |  |  |  |  |  |  | 4196195  T > G |
